# Design of combination therapeutics from protein response to drugs in ovarian cancer cells

**DOI:** 10.1101/2025.04.28.651139

**Authors:** Alexandra Franz, Ciyue Shen, Fabian Coscia, Kenneth Munroe, Lea Charaoui, Anil Korkut, Matthias Mann, Augustin Luna, Chris Sander

## Abstract

High-grade serous ovarian cancer (HGSOC) remains the most lethal gynecologic malignancy, and novel treatment approaches are needed. Here, we used unbiased quantitative protein mass spectrometry to assess the cellular response profile to drug perturbations in ovarian cancer cells for the rational design of potential combination therapies. Analysis of the perturbation profiles revealed proteins responding across several drug perturbations (called frequently responsive below) as well as drug-specific protein responses. The frequently responsive proteins included proteins that reflected general drug resistance mechanisms, such as changes in drug efflux pumps. Network analysis of drug-specific protein responses revealed known and potential novel markers of resistance, which were used to rationalize the design of anti-resistance drug pairs. We experimentally tested the anti-proliferative effects of 12 of the proposed drug combinations in 6 HGSOC cell lines. While response typically varies across different cell lines, 10 of the 12 combinations tested have either an additive or synergistic CI index in at least one cell line and may therefore be plausible candidates for overcoming or preventing resistance to single agents. Serendipitously, we observed an unexpectedly strong 0.05-0.11 micromolar response to GPX4 inhibitors as single agents in the OVCAR-4 cell line. We propose several drug combinations as potential therapeutic candidates in ovarian cancer, as well as GPX4 inhibitors as single agents.

**Highlights:**

- 5000-protein response to 7 drugs in an ovarian cancer cell line profiled by protein mass spectrometry
- Network analysis suggested potential pathways of drug resistance inferred from response profiles
- Demonstration of a general method for profiling adaptive response to therapeutic interventions with implications for the development of anti-resistance therapy
- Plausible anti-resistance drug combinations tested for antiproliferative effect in up to 6 ovarian cancer cell lines
- Drug combinations with additive or synergistic antiproliferative effects are plausible pre-clinical candidates for overcoming or preventing resistance to single agents
- Several combinations were synergistic in some cell lines (with PARPi, MEKi, and SRCi)
- We observed 0.05-0.11 micromolar response to GPX4 inhibitors as single agents in the OVCAR-4 cell line

## Introduction

### High-grade serous ovarian cancer has an unmet clinical need

High-grade serous ovarian cancer (HGSOC) is the most common and most aggressive subtype of ovarian cancer with a high mortality rate (Matulonis et al. 2016; Siegel, Miller, and Jemal 2019). Despite surgical removal and extensive chemotherapy as first-line treatments, approximately 80% of advanced-stage tumors relapse, resulting in a 5-year survival rate of less than 30% (Colombo, Lorusso, and Scollo 2017). Only small improvements in the overall survival rate have been seen in HGSOC patients over the past decades, carrying a grim prognosis (Berns and Bowtell 2012; Bowtell et al. 2015). There is thus a major unmet need for effective antitumor strategies to improve patient outcomes.

### Development approaches to drug combinations in cancer

One approach to developing effective treatment regimens is the deep characterization of cancer cellular response profiles to drug perturbations. This information has not only the potential to provide predictors of therapy response but also to determine optimal drug combinations (Al-Lazikani, Banerji, and Workman 2012). For example, one combination strategy is the co-administration of drugs with similar mechanisms of action (MoAs) to enhance target coverage and inactivation of oncogenic signaling. Successful examples are pertuzumab and trastuzumab to treat HER2^amp^ metastatic breast cancer (Baselga et al. 2012) and the combination of dabrafenib and trametinib to treat BRAF^V600E^ melanoma (Long et al. 2016; Robert et al. 2015). Another combination strategy, which we aim for here, is to block emerging resistance pathways that are activated in response to drug treatment and plausibly represent an evasive response to perturbation. For example, compensatory activation of MEK and/or AKT has been shown to limit mTOR/PI3K inhibitor therapy in prostate and breast cancer cell lines and patient tumor samples, which could be reversed by combined inhibition of mTOR/PI3K and AKT (Carracedo et al. 2008; O’Reilly et al. 2006). Other examples of drug pairs that work through parallel pathways and lead to enhanced anti-tumor efficacy include the combined inhibition of HER2 and PI3K in breast cancer cells (Rexer and Arteaga 2012) or co-targeting the HER family and IGF-R with afatinib and NVP-AEW541 in pancreatic cancer cells (Ioannou et al. 2013).

### Available post-perturbation drug response datasets

Over the last few years, there have been extensive efforts to decipher the mechanisms involved in drug response, including large-scale collections of gene expression response profiles across many cancer cell lines following drug exposure. This transcriptomics approach was pioneered by the Con-Map (Lamb et al. 2006) and extended by the NIH LINCS program (Koleti et al. 2018) and the L1000 project (Niepel et al. 2017). Considering that small molecule inhibitors usually target proteins, which are the basic functional units in biological processes, efforts have been made to collect proteomic response profiles in order to provide a more functional and proximal readout of drug action (R. F. S. Lee et al. 2017; Saei et al. 2019; Ruprecht et al. 2020; Yan et al. 2022). However, these efforts have been limited in ovarian cancer. So far, the biggest collections of drug response protein profiles in ovarian cancer cells cover 1) targeted antibody signal measurements of a total of 210 proteins and phosphorylation sites by antibody-based reverse-phase protein array (RPPA) in several ovarian cancer cell lines and 2) a study focused on BET bromodomain inhibition conducted in cell lines using mass spectrometry (Zhao et al. 2020; Gonçalves et al. 2022; Kurimchak et al. 2016). Unbiased large-scale proteomic approaches are still lacking.

### Identifying anti-resistance drug combinations from mass spectrometry-based proteomic profiles after drug treatment

To increase the scale of proteomic response profiles for ovarian cancer, we used unbiased deep mass spectrometry (MS)-based proteomics and comprehensively assessed the proteomic changes induced by drug action in order to identify candidate markers of resistance and provide information for drug combination strategies in ovarian cancer. We measured ovarian cancer cells’ proteomic response to a number of different anti-cancer drugs (STable 2) by protein mass spectrometry. Comparative analysis revealed specific protein responses for each of the drugs and general protein responses shared across several drugs. While we found that many generally responsive proteins in this dataset are involved in general drug resistance mechanisms (e.g., drug efflux pump), the data-driven protein network analysis of drug-specific protein responses suggested novel potential indicators of resistance response as well as known resistance indicators. Based on these observations, we propose several drug combinations for follow-up studies in ovarian cancer cells as an example of using comprehensive proteomic response profiling for the rational selection of drug combination therapeutic candidates.

## Results

### Computational results from proteomic response analysis to nominate drug combinations

#### Mass-spectrometry protein response profiling of anti-cancer compounds in ovarian cancer cells

First, we selected several anti-cancer drugs that are currently under development for the treatment of ovarian cancer patients. The selected drugs, MK-2206, Venetoclax, CHIR-99021, PD-0325901, Bisindolylmaleimide VIII (BIM VIII), and Bosutinib were selected similarly to a PARP inhibitor in our earlier study (Franz et al. 2021). Each drug targets a frequently activated oncogenic signaling pathway or biological process, such as the Akt (protein kinase B) pathway, apoptosis involving Bcl-2 (B-cell lymphoma 2), the GSK3 (glycogen synthase kinase 3) pathway, the MAPK/ERK pathway (mitogen-activated protein kinase cascade), PKC (protein kinase C) signaling, and SRC (protein tyrosine kinase) signaling. We used OVSAHO cells as a preclinical HGSOC model system, as this cell line has similar genomic and gene expression features as those measured in surgical HGSOC samples (Domcke et al. 2013; Coscia et al. 2016; Sinha et al. 2021). To explore the effects of these drugs on cellular protein levels, we perturbed OVSAHO cells with these inhibitors at their IC50 concentration (half-maximal inhibitory concentration) and measured the protein changes using MS. The IC50 for each drug was determined based on the dose-response curves derived from cell viability measured 72 hrs after drug treatment. The treatment time of 72 hrs is plausibly sufficient to capture the adaptive proteomic response of cells upon drug treatment (Arif, Datar, and Kalsotra 2017). Cell viability was measured by cell counting using live-cell imaging (Materials and Methods). Treatment with individual inhibitors resulted in a dose-dependent reduction in cell viability (Figure 1a) with IC50 values ranging from 4 µM to 12 µM (STable1). OVSAHO cells were treated at each drug’s IC50 concentrations for 72 hrs in three biological replicates, and proteomic changes were profiled by MS (Figure 1b). To quantify relative protein levels across drug perturbation conditions, we applied data-independent acquisition (DIA) combined with label-free based quantification (Materials and Methods). We quantified drug-perturbed protein profiles at a median depth of ∼5,000 protein groups (groups are indistinguishable based on peptides identified) per perturbation condition (Figure 1c). Unsupervised multi-dimensional scaling (MDS, Euclidean distance) of biological replicates confirmed reasonable reproducibility of proteomic responses upon each drug treatment (Figure 1d).

**Figure 1.**
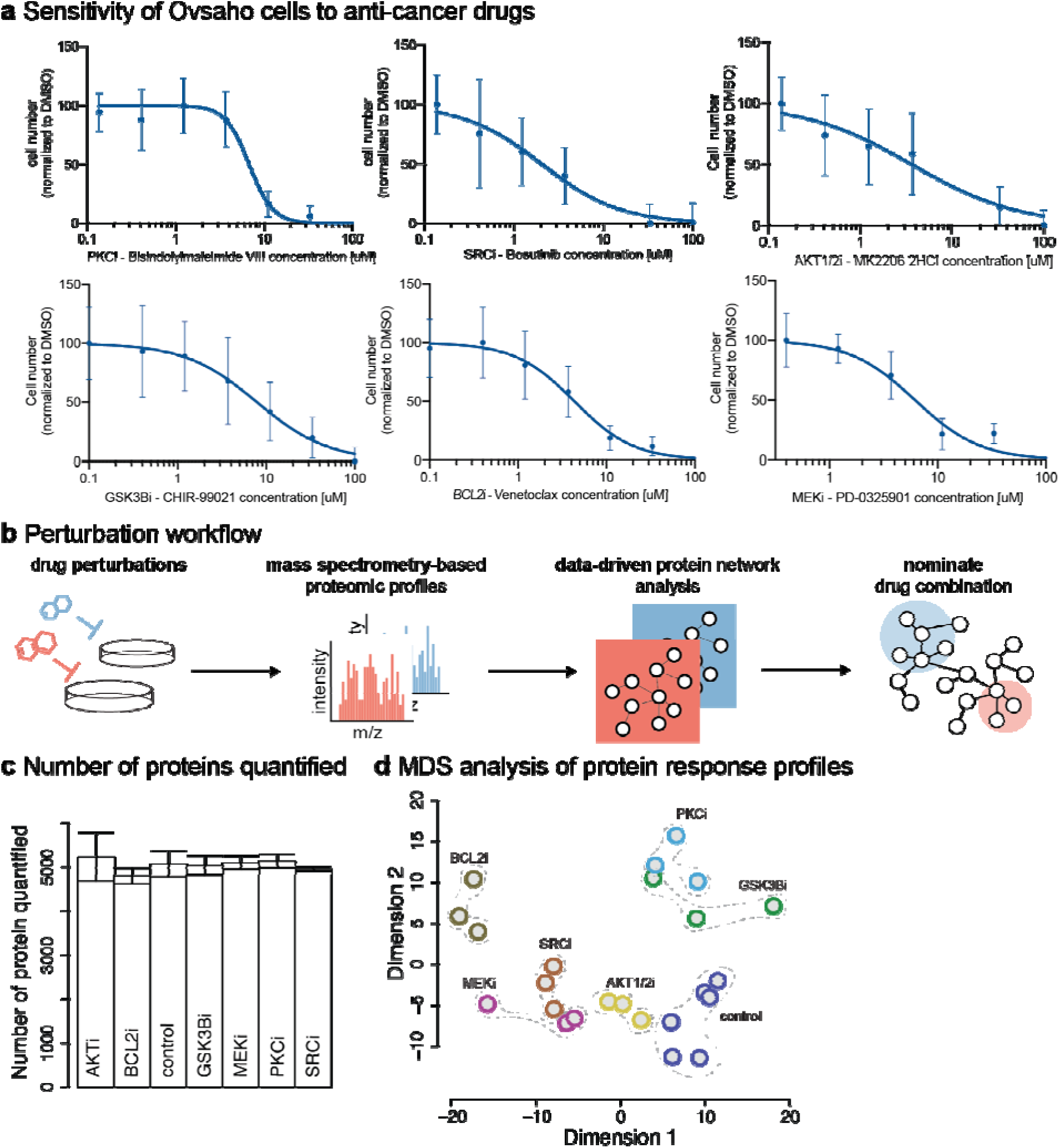
Acquisition of proteomic changes following molecularly targeted drug treatments in OVSAHO cells. (a) Dose-response curves of the single small-molecule inhibitors, used for the primary perturbation followed by MS protein response profiling in OVSAHO cells, including AKT inhibitor MK-2206 (AKTi), BCL2 inhibitor Venetoclax (BCL2i), GSK3β inhibitor CHIR-99021 (GSK3βi), MEK inhibitor PD-0325901 (MEKi), PKC inhibitor Bisindolylmaleimide VIII (BIM VIII, PKCi), and SRC inhibitor Bosutinib (SRCi). Data is aggregated from three biological replicates, each with three technical replicates, and error bars represent standard deviations of the data. (b) Label-free mass spectrometry (MS)-based proteomics workflow used to measure protein changes in response to drug treatment at IC50. Data-driven network analysis was later applied to the acquired MS data to identify potential resistance mechanisms for the nomination of drug combination candidates. (c) Number of proteins quantified upon each drug treatment of OVSAHO cells after 72h at inhibitor IC50 concentration. Error bars: standard deviation of three biological replicates for each condition. (d) Multidimensional scaling (MDS, Euclidean distance) 2D projection of protein levels measured after treatment. Colors indicate different drug perturbation conditions with weak dotted lines around each group of biological replicates to indicate the closeness of the replicates.

#### Comparison of strongly responsive proteins across drug perturbation conditions

We examined the measured global proteome response for each of the six drugs and identified a total of 5858 proteins, of which 4480 (76%) proteins were experimentally detected in all perturbation conditions (SFigure 1). We assessed which proteins and cellular processes were most strongly affected by the drug treatment. We defined strongly responsive proteins as proteins with an expression level change upon drug treatment compared to the control treatment of at least 0.5 or -0.5 (log2 ratio of perturbed/control) with a nominal p-value < 0.05 and BH-based FDR < 0.2 (Materials and Methods). Using these parameters, we found that most proteins (>80%) did not respond strongly across treatment conditions (Figure 2a). The different drug treatments resulted in different proportions of strongly responsive proteins, which indicates the different extent and scope of downstream signaling. For example, while AKTi induced strong expression level changes in a relatively small number of proteins (8% of the measured proteins), inhibition of SRC affected as many as 18%. Indeed, SRC activity is known to have key roles in affecting multiple pathways (Mayer and Krop 2010). Across the six different perturbations, the expression levels of a total of 2714 proteins changed significantly in at least one drug condition (‘strong response’), of which only 127 proteins changed strongly in at least 4 perturbation conditions (‘frequently responsive’) (Figure 2b; STable2). Frequently affected proteins generally had expression level changes in the same direction across all drug perturbations (Figure 2c). 115 out of 127 proteins (91%) have consistent directional changes, indicating a potential general stress response. The strongly responsive proteins for each drug are used below as a guide for choosing candidates for anti-resistance drug combinations.

**Figure 2.**
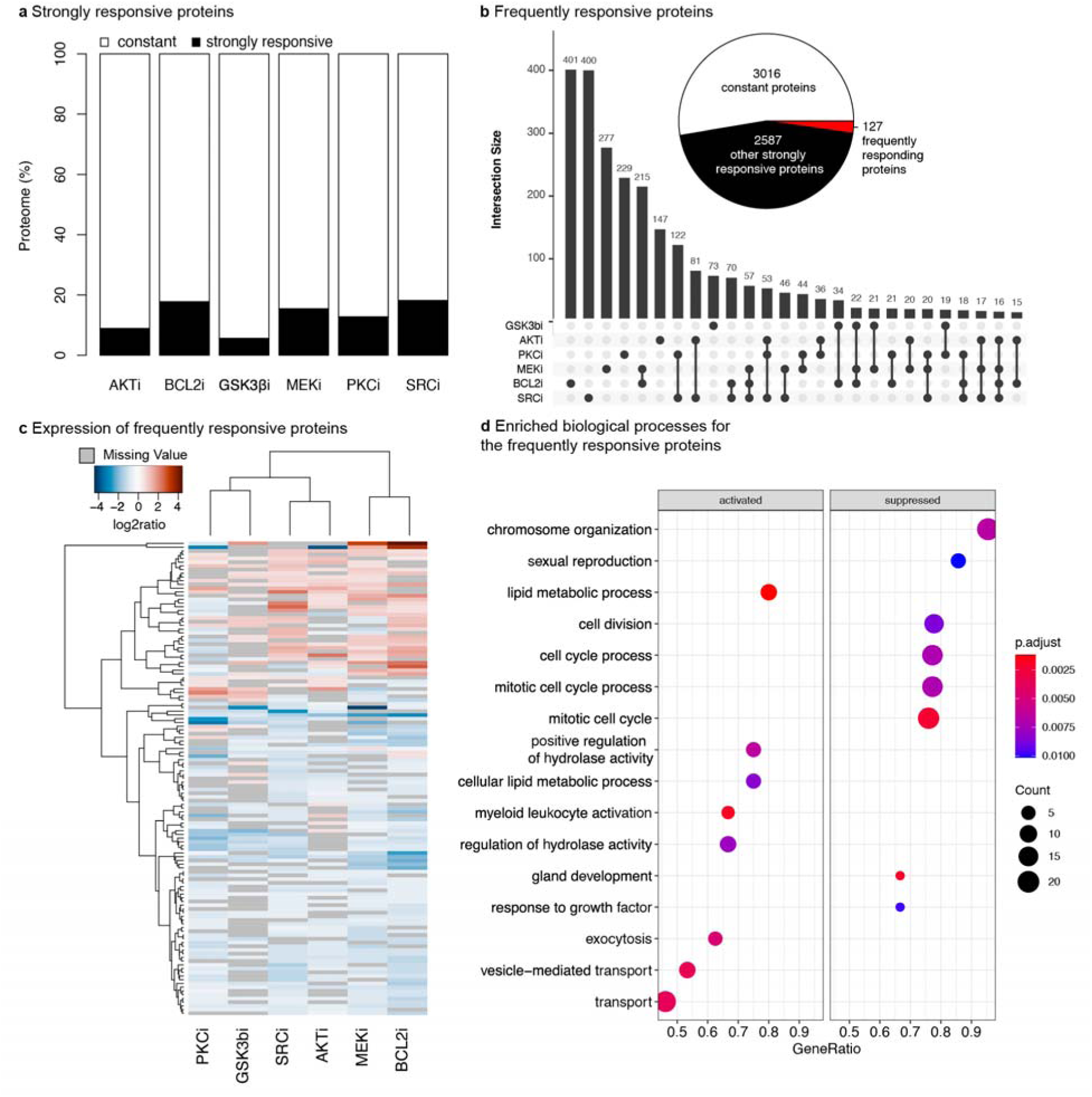
Identification of frequently responsive proteins and processes induced by pharmacological inhibitors. **(a)** Proportion of proteins per treatment condition that changed compared to the control treatment. Strongly responsive proteins are defined as proteins whose absolute log2 expression change is at least 0.5 (p-value < 0.05 and BH-based FDR < 0.2 in t-test). Constant proteins are the remaining proteins; protein expression values are averaged over three biological replicates. **(b)** Number of proteins that strongly respond to one or more drugs are shown in the upset plot. Intersection sets with more than 15 proteins are shown. Proteins that strongly respond in 4 or more treatment conditions are defined as frequently responsive proteins (red). The set of 2587 drug-specific proteins (black in the pie chart) is the union of the strongly responsive proteins for the six drugs (black rectangles in (a)). **(c)** Protein expression change of identified frequently responsive proteins; “frequently” refers to both positively and negatively responding proteins in at least 4 conditions (red in (b)). Most proteins (115/127) respond with consistent trends across perturbation conditions (defined as the directions of change are consistent in all but at most one drug), indicating a general stress response. **(d)** Gene set enrichment analysis (GSEA) identified both activated (increased protein expression) and suppressed (decreased protein expression) biological processes. Suppressed processes likely reflect the cytotoxic effects of drugs, while activated processes might suggest a general resistance mechanism.

#### Identification of general response markers to pharmacological inhibition

Several well-known resistance mechanisms have been observed to develop in cancer cells against different pharmacological compounds that are structurally and functionally unrelated (Zelcer et al. 2001). Therefore, we next assessed the functional properties of the 127 frequently responsive proteins. Using gene set enrichment analysis (GSEA) (Subramanian et al. 2005), we found that levels of proteins involved in cell survival and proliferation were significantly decreased, including in biological processes such as chromosome organization (GO:0051276, p-value = 6.63E-03), and cell cycle processes (GO:0022402, p-value = 7.18E-03) (Figure 2d). In contrast, biological processes, such as lipid metabolic processes (GO:0006629, p-value = 1.19E-03), vesicle-mediated transport (GO:0016192, p-value = 3.54E-03), and positive regulation of hydrolase activity (GO:0051345, p-value = 6.43E-03), included proteins with significantly increased expression as a consequence of drug perturbation. Consistent with previous research on the response of cancer cells under stressed environments (Munir et al. 2019; Q. Wu et al. 2021), these up-regulated processes are likely general adaptive responses of ovarian cancer cells to pharmacological inhibition.

#### Identification of drug-specific response markers and nomination of drug combinations

We next analyzed strongly responsive proteins for each individual drug treatment condition to gain further insight into drug-specific cellular response. We applied three independent and complementary computational approaches to nominate drug-responsive proteins and pathways. The approaches include the NetBox algorithm for module detection (Cerami et al. 2010; Liu et al. 2020), gene set enrichment analysis (GSEA) (Subramanian et al. 2005), and interpretation of individual protein expression changes by inspection of functional annotations. The NetBox algorithm combines prior knowledge of protein interaction networks with a clustering algorithm to identify functional protein modules. We used the NetBox algorithm to identify likely functional protein modules and used annotations of individual protein functions of module members to functionally label each module (Materials and Methods). We applied GSEA to independently identify affected biological processes by each drug inhibition. In addition, we examined individual responsive proteins that are reported to have a role in known cancer signaling. For each of the small-molecule drugs, the functional annotation of the identified protein modules with decreased activities generally agreed with the known inhibitory roles of the drugs (Figure 3a, Supplementary notes, SFigure 2). The identified modules and their protein members are of particular interest as they are likely indicative of potential adaptation or resistance mechanisms to the small-molecule inhibitors.

**Figure 3.**
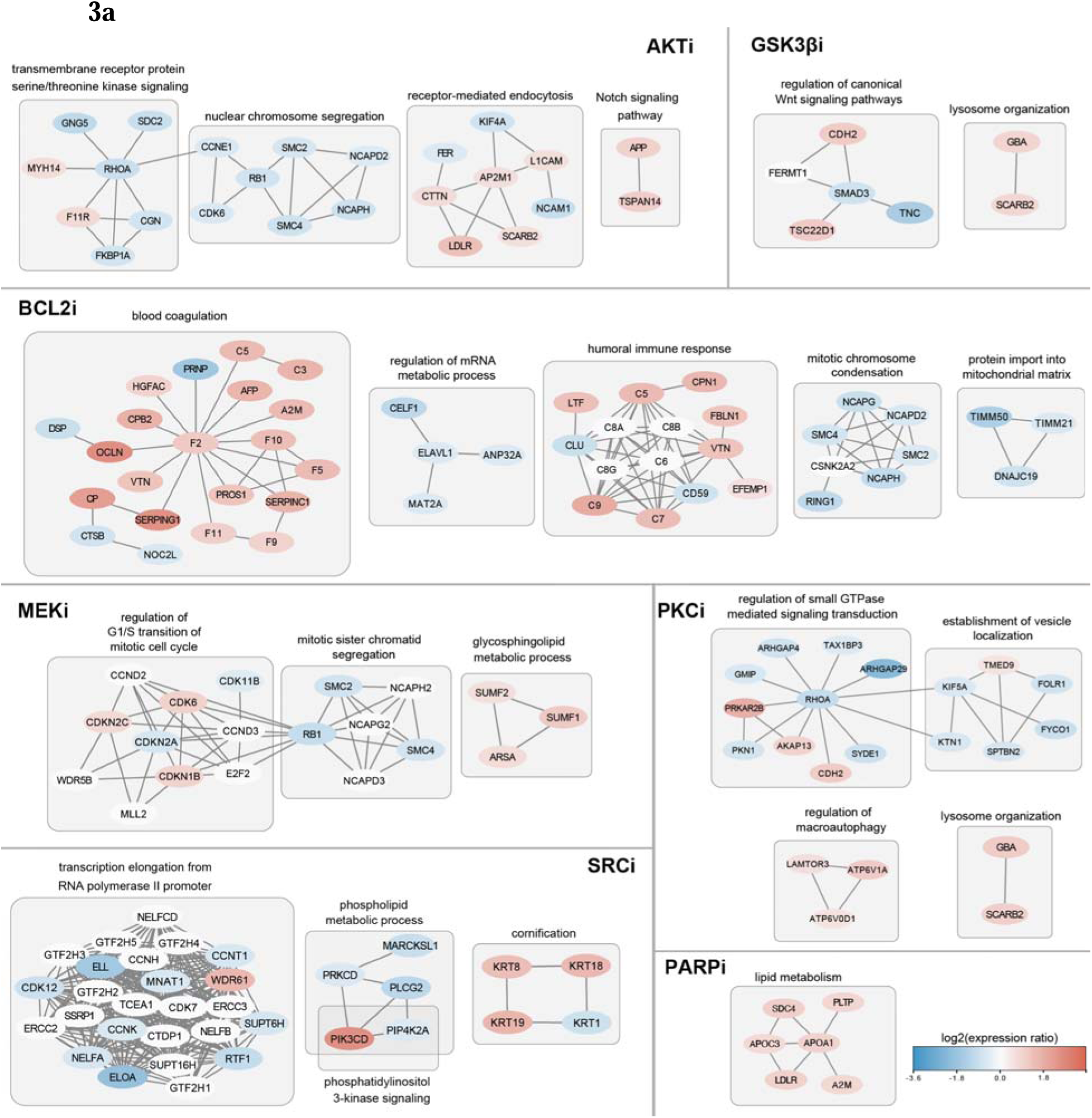
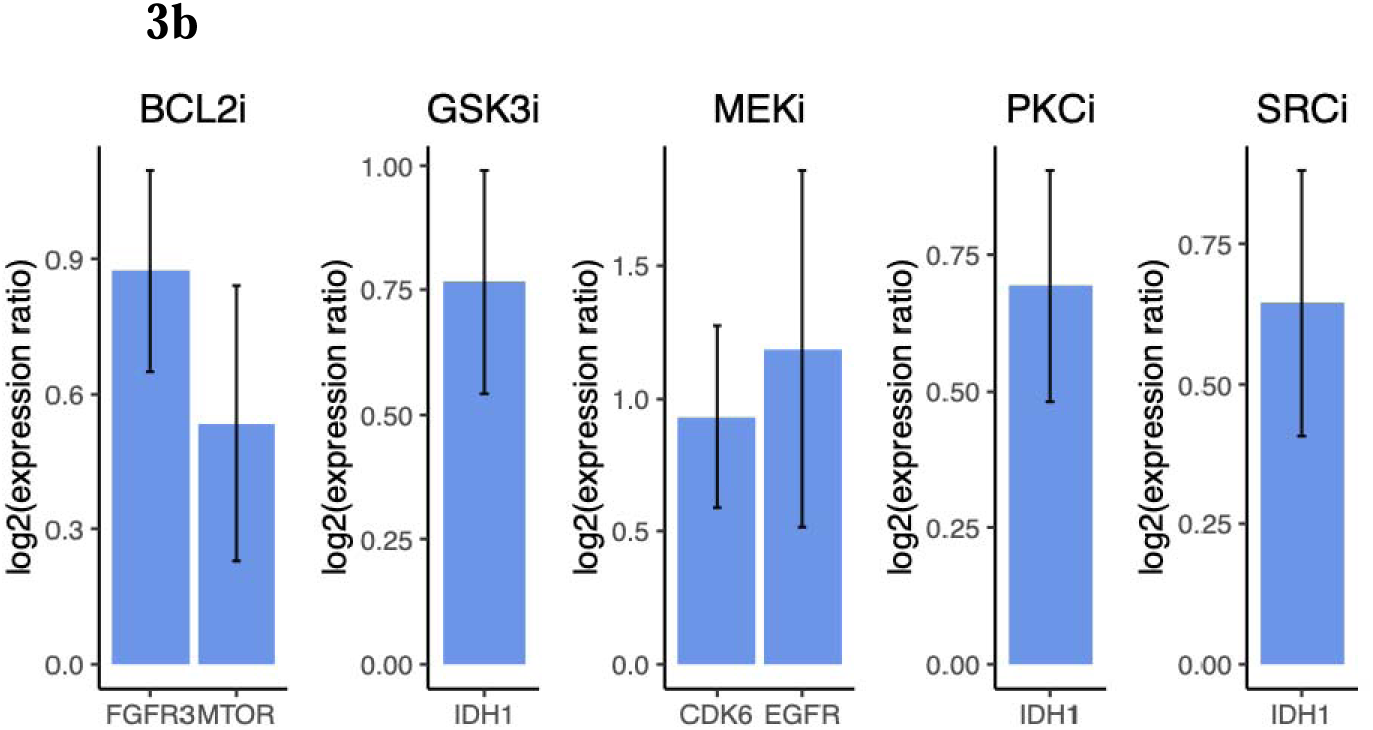
Drug-specific protein response networks identified using NetBox. (**a**) Drug-specific protein responses were grouped into functional modules using the NetBox algorithm, and the modules were characterized by functional labels based on enrichment analysis (Materials and Methods). Subsets of protein modules for each drug perturbation were chosen based on their relevance to the known functions of the inhibitors and their implications for potential resistance mechanisms. Nodes are protein colored by protein expression change, log2(expression ratio perturbed/unperturbed). Edges are undirected protein interactions from the background network used by the NetBox algorithm (Reactome FI Network or INDRA network). (**b**) Individual protein expression change ratios that are justification for the nominated drug combinations.

We identified potential resistance mechanisms as protein modules and biological processes that are pro-proliferative and increased in activity, as indicated by an increase in average protein expression, or anti-proliferative and decreased in activity. Whether a given protein or a biological process is pro-proliferative or anti-proliferative is defined as an individual functional score (1: pro-proliferative or -1: anti-proliferative) for the protein or the average functional score of the proteins in the process (Materials and Methods). To nominate potentially therapeutic drug combinations, we suggest targeting proteins involved in the presumed resistance mechanisms together with the original small-molecule target, with the intent to block the restoration of proliferation. The candidate drugs were chosen as specific inhibitors of the selected pathways (Table 1). We then ranked the drug combination candidates based on their clinical stages, prioritizing drugs approved by the U.S. Food and Drug Administration (FDA) (e.g., rucaparib, an approved treatment, versus AG14361, an experimental compound). To further filter th candidates, we also reviewed previous literature on how widely these candidates have been used in preclinical studies, as well as additional supporting information on resistance mechanisms and potential drug pairs in combination therapies (see Supplementary notes).

**Table 1.**
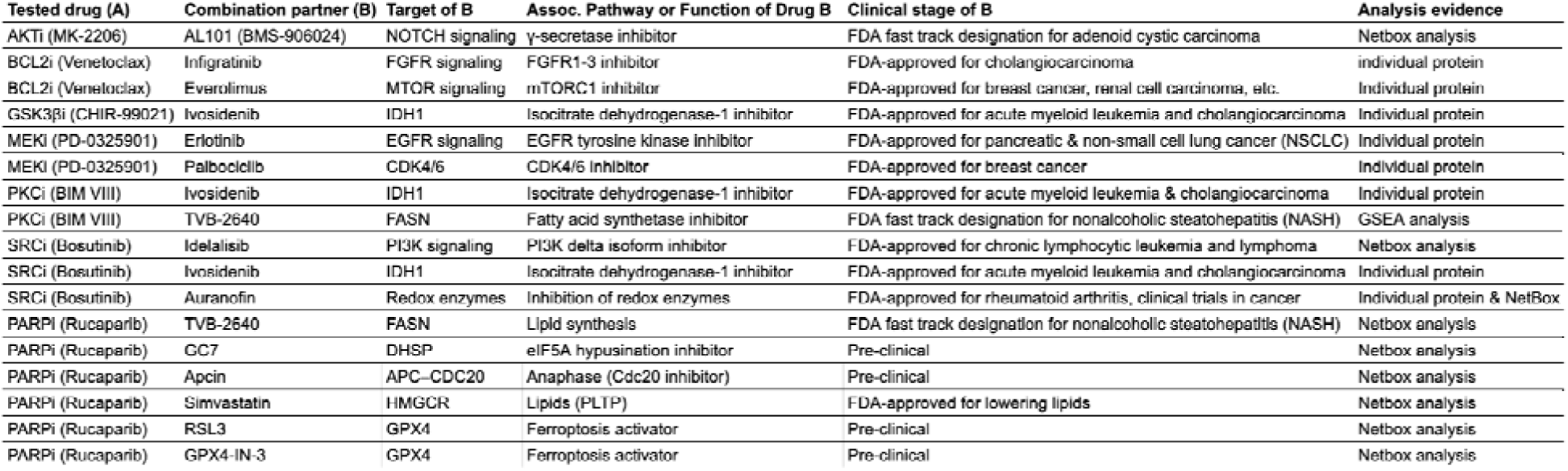
Proposed combination drug candidates based on the analysis of the proteomic response profiles. These combinations were tested experimentally. Potential resistance mechanisms were identified based on the analysis of proteomic profiling after drug treatment using three complementary approaches (i.e., NetBox analysis, GSEA analysis, and individual protein expression analysis). Small molecule inhibitors were selected to target the corresponding pathways indicative of resistance as combination partners of the originally profiled drugs. Drugs are prioritized if they are more relevant in clinical settings, i.e., FDA-approved or in late-stage clinical trials.

### Potential drug combination candidates with PARPi

Identifying drug pairs for combination intervention is meant to address the problem of lack of sensitivity to PARPi in some ovarian tumors in a clinical setting, an active area of research (Dréan, Lord, and Ashworth 2016). Our computational results indicate that an inferred module of interest with increased protein expression after perturbation with the PARPi olaparib in OVSAHO cells contains a set of proteins involved in lipid metabolism and lipoprotein processe and reported to pairwise interact (Figure 3a): A2M (alpha 2 macroglobulin, interacts with APO lipoproteins), SDC4 (syndecan4, a plasma membrane proteoglycan), PLTP (a phospholipid transfer protein, involved in the transfer of excess surface lipids from triglyceride-rich lipoproteins to HDL and involved in the uptake of cholesterol from peripheral cells and tissues), LDLR (a member of the low-density lipoprotein receptors family), APOC3 (reported to promote the assembly and secretion of triglyceride-rich VLDL particles) and ApoA1 (a protein component of high-density lipoprotein particles).

As the abundance of each of these five functionally connected proteins increased after PARPi perturbation, we searched for drugs that affect cellular processes involving lipids or lipoproteins by targeting these proteins directly or indirectly. After inspection of pathway knowledge bases and background knowledge of molecular processes involved in resistance to PARP inhibition, we chose the following drugs to be combined with PARPi. (1) TVB, a fatty acid synthase (FASN) inhibitor, which reduces fatty acid turnover. (2) RSL3 and GPX4-IN-3, both inhibitors of GPX4. GPX4 is a selenoprotein that reduces lipid hydroperoxides and prevents ferroptosis in conditions of oxidative stress (ROS) with enhanced lipid peroxidation, to the advantage of surviving cancer cells (Yang et al. 2014). Ferroptosis is a form of regulated cell death characterized by the accumulation of lipid peroxides; so, inhibiting GPX4 promotes ferroptosis, reducing cell count. (3) Simvastatin, an inhibitor of HMG-CoA reductase (HMGR), is an enzyme in the mevalonate pathway and the rate-limiting enzyme in cholesterol biosynthesis; the reason for inclusion is that statins are reported to have many pleiotropic effects downstream of the mevalonate pathway and to have anti-invasive and anti-inflammatory effects supporting a repurposed anti-cancer use (Matusewicz et al. 2015) (Zaky et al. 2023). In addition, as motivated in our previous report (Franz et al. 2021), we also added the drugs (4) GC7, inhibitor of deoxyhypusine synthase, and (5) apcin, inhibitor of the activity of the anaphase-promoting complex APC/C, to be combined with PARP inhibition (Table 1). An additional proposed combination involves rucaparib with novobiocin, an inhibitor of polymerase theta, motivated independently (not as a result of our proteomics screen) due to interest in the work of d’Andrea et al. (Iorio et al. 2016; Zhou et al. 2021), who reported that BRCA-deficient tumor cells with acquired resistance to PARPi are sensitive to novobiocin in vitro and in vivo.

### Potential drug combination candidates with AKTi

We observed that upon AKT inhibition, a NetBox-identified protein module enriched for the Notch signaling pathway (GO:0007219, adj. p-value = 2.37E-02) has increased expression levels with an average log2 expression ratio (after/before perturbation) of 0.99 over two genes, APP and TSPAN14. Activation of Notch signaling is found in different types of cancer, and inhibition of Notch signaling has been shown as a potential therapeutic approach (Katoh and Katoh 2020). Thus, the upregulation of proteins in Notch signaling pathways might be an adaptive response for ovarian cancer cells to resist AKT inhibition. Therefore, we propose to use AL101 (BMS-906024), a γ-secretase inhibitor, which blocks Notch activity by preventing cleavage of the Notch receptors at the cell surface (Olsauskas-Kuprys, Zlobin, and Osipo 2013) in combination with the AKT inhibitor.

### Potential drug combination candidates with BCL2i

We identified two pro-proliferative genes whose protein expression increased upon BCL2 inhibition, FGFR3 (log2ratio = 0.87, p-value = 2.00E-03) and MTOR (log2ratio = 0.53, p-value = 3.49E-02), which have well-known roles in cancer signaling, i.e., FGFR signaling and mTOR signaling (Figure 3b). With additional evidence from previous literature reporting enhanced anti-tumor effects of dual inhibition (inhibition of Bcl-2 and FGFR in endometrial cancer, and of Bcl-2 and mTOR in renal cell carcinoma) (Packer et al. 2019; Nayman et al. 2019), we hypothesize that both signaling pathways may be able to implement resistance to BCL2 inhibition in OVCA and may contain reasonable combination targets with BCL2i. We chose Infigratinib, a selective FGFR1-3 inhibitor to target FGFR signaling (FDA-approved for treatment of cholangiocarcinoma with an FGFR2 fusion), and Everolimus which inhibits the formation of the mTOR complex, for testing in combination with BCL2i. Everolimus has been the subject of Phase I and II clinical trials testing certain drug combinations in ovarian cancer (NCT01281514, NCT00886691).

### Potential drug combination candidates with GSK3***β***i

We found that the protein expression of isocitrate dehydrogenase 1 (IDH1) increased upon GSK3β inhibition (log2ratio = 0.767, p-value = 1.40E-03) (Figure 3b). It has been previously reported that pharmacological inhibition or knockdown of IDH1 decreased proliferation by inducing senescence in multiple HGSOC cell lines (Dahl et al. 2019). Therefore, we believe IDH1 can be a resistance marker for GSK3β inhibition. We select Ivosidenib, an FDA-approved inhibitor for IDH1, as the combination partner for GSK3βi.

We also found that the NetBox-identified protein module enriched for lysosome organization (GO:0007040, adj. p-value = 1.18E-03) is activated. This response has been previously reported by Albrecht et al. (Albrecht et al. 2020), who observed that macropinocytosis, triggered by GSK3-inhibition, induced catabolic activity of lysosomes. We believe increased lysosome activity might be a resistance response to GSK3β inhibition. The lysosome is linked to many hallmarks of cancer, and a wide range of agents have been shown to affect multiple aspects of lysosome activities in clinical trials (Davidson and Vander Heiden 2017). We believe our analysis does not provide sufficient evidence to support our choice of drug(s) inhibiting a particular lysosomal target. Therefore, experimental validation of co-targeting GSK3β and lysosome activities is beyond the scope of this study.

### Potential drug combination candidates with MEKi

The protein expression of EGFR (log2ratio = 1.19, p-value = 1.01E-02) increased upon MEK inhibition (Figure 3b). Previous literature demonstrated that combined inhibition of MEK and EGFR prevents the emergence of resistance in EGFR-mutant lung cancer. We, therefore, selected the most widely used EGFR inhibitor, Erlotinib, as the combination partner with our MEKi. We also observed that the expression of cyclin-dependent kinase 6 (CDK6) increased in response to MEK inhibition (log2ratio = 0.931, p-value = 4.55E-03), which positively affects the cell cycle. Since CDK6 is a potential resistance marker of MEKi, as well as a single therapeutic target for ovarian cancer patients (Dall’Acqua et al. 2021), we selected palbociclib as the specific inhibitor of CDK4/6 to use in combination with MEKi.

We found an increase in the expression of proteins enriched for the glycosphingolipid metabolic process (GO:0006687, adj. p-value = 3.51E-07) using NetBox. Interestingly, metabolic rewiring in response to MEK inhibition has been previously described as a possible mechanism of resistance in melanoma cells (Ruocco et al. 2019; Nguyen et al. 2020), and targeting of lipid metabolism has been previously reported in ovarian cancer cells (Chen et al. 2019). Although targeting highly expressed metabolic enzymes in the glycosphingolipid metabolic process might provide a potential strategy to overcome MEKi resistance in ovarian cancer, this hypothesis has yet to be tested due to the lack of specific inhibitors of the pathway.

### Potential drug combination candidates with PKCi

The results of GSEA suggested that proteins involved in lipid metabolic process (GO:0006629, p-value = 1.13E-03) have increased expression upon PKC inhibition. Lipid metabolic processes have been identified as part of the general stress response and as potential targets for combination therapy (Chen et al. 2019; Snaebjornsson, Janaki-Raman, and Schulze 2020). Therefore, we selected TVB-2640, a fatty acid synthase inhibitor, to target this potential resistance mechanism. Similar to our observations with GSK3β inhibition, IDH1 (log2ratio = 0.694, p-value = 1.81E-03), as well as proteins involved in the process of lysosome organization (GO:0007040, adj. p-value = 1.24E-05), have increased expression upon PKCi (Figure 3b). We propose Ivosidenib with PKCi as a potential combination candidate.

### Potential drug combination candidates with SRCi

We found that one responsive protein module identified by NetBox is enriched for proteins involved in the phospholipid metabolic process (GO:0006644, adj. p-value = 4.86E-04). An interesting observation is that two proteins, PI3K delta isoform (PIK3CD) and phosphatidylinositol-5-phosphate 4-kinase type-2 alpha (PIP4K2A) in this module are also involved in the PI3K signaling pathway, which is critical for cell survival and frequently altered in ovarian cancer. Although SRC inhibition has been previously reported to suppress the PI3K pathway (Beadnell et al. 2018), our observation suggested that the protein expression of the PI3K delta isoform significantly increased upon SRC inhibition (log2ratio = 2.48, p-value = 1.77E-03). Therefore, we propose to use Idelalisib, a specific PI3K delta isoform inhibitor, in combination with SRCi. The combination of SRC and PI3K (by saracatinib and GDC-0941, respectively) has been shown to be effective in renal cancer in several pre-clinical models, including PDX models (Roelants et al. 2018). We also observed an increase in protein expression of IDH1 upon SRC inhibition (log2ratio = 0.644, p-value = 2.76E-03) and therefore propose Ivosidenib and SRCi as a combination candidate (Figure 3b). In addition, we observed that the protein expressions of peroxiredoxins II (log2ratio = 0.754, p-value = 2.87E-03) and V (log2ratio = 0.821, p-value = 8.95E-04) (PRDX) are significantly increased. PRDXs are antioxidant enzymes that play key roles in regulating peroxide levels within cells, controlling various physiological functions (Perkins et al. 2015). Several studies have implicated an increase in reactive oxygen species (ROS) in carcinogenesis due to a loss of proper redox control (Nicolussi et al. 2017). Most notably, in ovarian cancer cells, PRDX expression was found to be associated with platinum drug resistance (X.-Y. Wang, Wang, and Li 2013) since increased levels of antioxidants inhibit apoptosis (Kalinina et al. 2012). Our NetBox analysis with strongly responsive protein identified a module labeled detoxification of ROS (data not shown). Our observations, together with the reported results of others, suggest that PRDXs are promising resistance markers to b targeted together with SRCi. Therefore, to target peroxiredoxins and the antioxidation process, we select auranofin, which inhibits the redox enzymes. Although not specific, auranofin is the closest to clinical use in cancer compared to all other inhibitors of antioxidation (Bajor et al. 2020).

### Experimental results on nominated drug combinations

#### Choice of cell lines for experimental validation

We experimentally tested the cellular response to drug combination candidates (Table 2, SFigure 3) proposed by our computational analysis of the protein perturbation response data (Table 1) that includes combinations based on our earlier work (Franz et al. 2021).

**Table 2.**
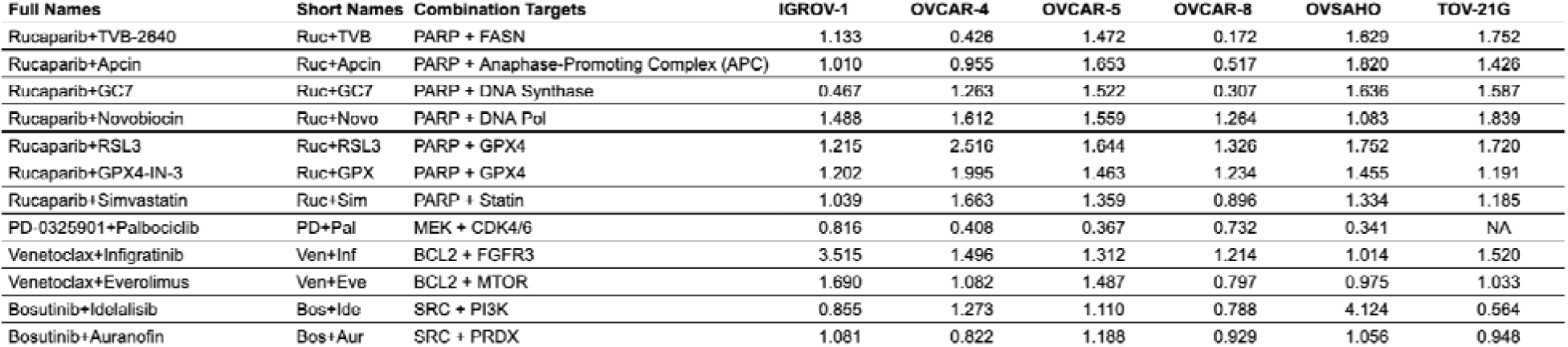
Drug combinations experimentally tested for effect on proliferation. The most interesting drug combinations were experimentally tested across several ovarian cell lines, and the inferred combination indices (CI) were calculated by the Chou-Talalay method. For drug combinations with 3 technical replicates, cell response was averaged across the replicates to perform the CI computation. IGROV-1, etc. (columns) are cell lines. NA: not available for technical reasons. All combinations with additive (0.9 < CI < 1.1) or synergistic (CI < 0.9) antiproliferative effects are considered potentially useful for further preclinical investigation, while clearly antagonistic ones (CI > 3.3) are not (Chou 2011).

We first tested each of the drug combinations in the OVSAHO cell line. For selected candidate combinations, we further tested the general validity of the anti-proliferative effects in another 5 HGSOC cell lines covering diverse genetic backgrounds (IGROV-1, OVCAR-4, OVCAR-5, OVCAR-8, and TOV-21G) (Table 2). These cell lines reflect a range of genomic and proteomic features of HGSOC patient tumors (Table 3). With respect to rucaparib, the cell lines used here cover a range of sensitivities. They have IC-50s for the PARPi rucaparib in the range of 13 to 206 uM as reported in the Genomics of Drug Sensitivity in Cancer (GDSC2, v8.5) dataset (Iorio et al. 2016), with OVCAR-4 and OVCAR-5 being the least sensitive to PARP inhibition (the OVSAHO cell line is not in GDSC) (Table 3). Together with OVASHO, these cell lines reflect well-defined preclinical models of HGSOC (Domcke et al. 2013; Coscia et al. 2016; Sinha et al. 2021). For each of the drug combinations, we measured cell viability after 72 hours.

**Table 3.** OVCA HGSOC cell line measures of similarity to patient samples and drug response to rucaparib. Cell line similarity to TCGA HGSOC patient samples a characterized by Domcke et al. and TumorComparer (TC) (Sinha et al. 2021; Domcke et al. 2013). Additionally, a select set of genes, including ones commonly altered in HGSOC (i.e., BRCA1, BRCA2, RB1) (Cancer Genome Atlas Research Network 2011). Rucaparib IC50 (µM, half-maximal inhibitory concentration) single drug screening of the cell lines as reported in Genomics of Drug Sensitivity in Cancer (GDSC) (Iorio et al. 2016). Mutations are protein-coding mutations as measured in particular cell line studies; similarly, CNAs are copy number alterations. KRAS, etc. are proteins, altered (green check) or not (blank) by mutation or CNA.

| Cell Line | GDSC Rucaparib IC50 (μM) | TC Similarity to Tumors (Higher Better) | Domcke Similarity to Tumors Rank (47 Total; Smaller Better) | Replication Time | # Mutations (cBioPortal) | # CNAs (cBioPortal) | KRAS | BRCA1 | BRCA2 | ATM | ATR | PTEN | PIK3CA | RB1 | TP53 |
| --- | --- | --- | --- | --- | --- | --- | --- | --- | --- | --- | --- | --- | --- | --- | --- |
| IGROV-1 | 13.45 | 0.46 | 43 | 20-30 h | 273 | 360 |  | ✓ | ✓ | ✓ |  | ✓ | ✓ | ✓ | ✓ |
| OVCAR-4 | 116.08 | 0.99 | 5 | 34-41 h | 38 | 2355 |  |  | ✓ | ✓ | ✓ | ✓ |  |  | ✓ |
| OVCAR-5 | 208.32 | 0.46 | NA | 27-50 h | NA | NA | ✓ |  |  |  |  |  |  |  |  |
| OVCAR-8 | 36.98 | 0.7 | 26 | 24-32 h | 42 | 1349 | ✓ |  |  | ✓ |  |  |  |  | ✓ |
| TOV-21G | 35.88 | 0.13 | 47 | 27-36 h | 154 | 147 | ✓ |  |  |  |  | ✓ | ✓ |  |  |
| OVSAHO | NA | NA | 2 | 36-72 h | 32 | 1118 |  |  | ✓ |  |  |  |  | ✓ | ✓ |

#### Several combinations with PARPi are anti-resistance candidates in ovarian cancer cells

For clinical therapeutic applications, both additive and synergistic combination drug effects are of interest. On the one hand, synergy implies an initial effect that is stronger than additive, deemed to be beneficial since side effects decrease with lower drug concentration. On the other hand, if resistance develops to one of the drugs, synergy is also lost, which is disadvantageous. Additionally, it is possible to see the benefit of combination therapy due to variable response in the population, even without synergy or additivity (Palmer and Sorger 2017).

For the purpose of careful consideration for future pre-clinical investigation, we report synergy values for all drug combinations for which there is sufficiently clear data. To quantify drug synergy (Combination Index, CI), we used the Chou-Talalay method (Chou 2010). All combinations with additive (0.9 < CI < 1.1) or synergistic (CI < 0.9) antiproliferative effect are of potential interest for further preclinical investigation, while clearly antagonistic ones (CI > 3.3) are not (Chou 2011). We observe that some combinations of PARP inhibition via rucaparib with compounds targeting several other targets have synergistic effects (CI < 0.9) in 3 of the 6 HGSOC cell lines: with (i) a fatty acid synthase inhibitor (TVB-2640), (ii) an APC inhibitor (Apcin), and (iii) an eIF5A-2 inhibitor (GC7). For PARP+FASN inhibition, we observe synergistic effects in both OVCAR-4 and OVCAR-8. The synergistic effect of PARP+FASN inhibition is most striking in OVCAR-8: the antiproliferative effect of the combination is nearly 6 times stronger than that of the single drugs. Interestingly, the use of the FDA-approved rucaparib PARP inhibitor did not produce the same additive or synergistic effects as combinations (inhibitors of FASN, APC, DNA synthesis, proteasome) with an alternative experimental PARP inhibitor (AG-14361), presumably due to differences in drug action on PARP activity. Given the importance of PARPi in clinical practice, several combinations with PARPi are therefore reasonable candidates for anti-resistance therapy in ovarian cancer.

#### Effects of combinations targeting GPX4 in ovarian cancer cells

A separate category of combinations tested experimentally involved the protein GPX4. GPX4 dependencies can arise in some therapy-resistant cancers (Hangauer et al. 2017; X. Wu et al. 2022). Both RSL3 and GPX4-IN-3 are preclinical compounds. Neither of these combinations resulted in synergistic combinations as defined by the CI value. We did, however, observe that GPX4 inhibitors have a remarkably strong single-agent effect in the cell lines tested, in the OVCAR-4 cell line at an IC50 of 0.11 micromolar for GPX4-IN3 and 0.05 micromolar for RSL3 (Figure 4, STable 3). It has been reported that an R245W mutation in TP53 may produce a protective effect against RSL3 in triple-negative breast cancer (TNBC) (Dibra et al. 2024). However, OVCAR-4 has a different mutation in TP53 (L130V), and that does not appear to have a protective effect against GPX4 inhibition, given the strong response we observe. Separately, there have been previous reports of synergistic combination effects with the alternative PARP inhibitor, olaparib, with RSL3 in the HEY HGSOC cell line; HEY is a G12D KRAS-mutated line (Hong et al. 2021). The authors of the previous study suggest that this combination works due to an adaptive response whereby GPX4 is induced in response to olaparib treatment. However, the sensitivity to single-agent GPX4 inhibition may be more general and have other explanations (Lei et al. 2024); detailed exploration of the mechanism of GPX4i is beyond the scope of the current study.

**Figure 4.**
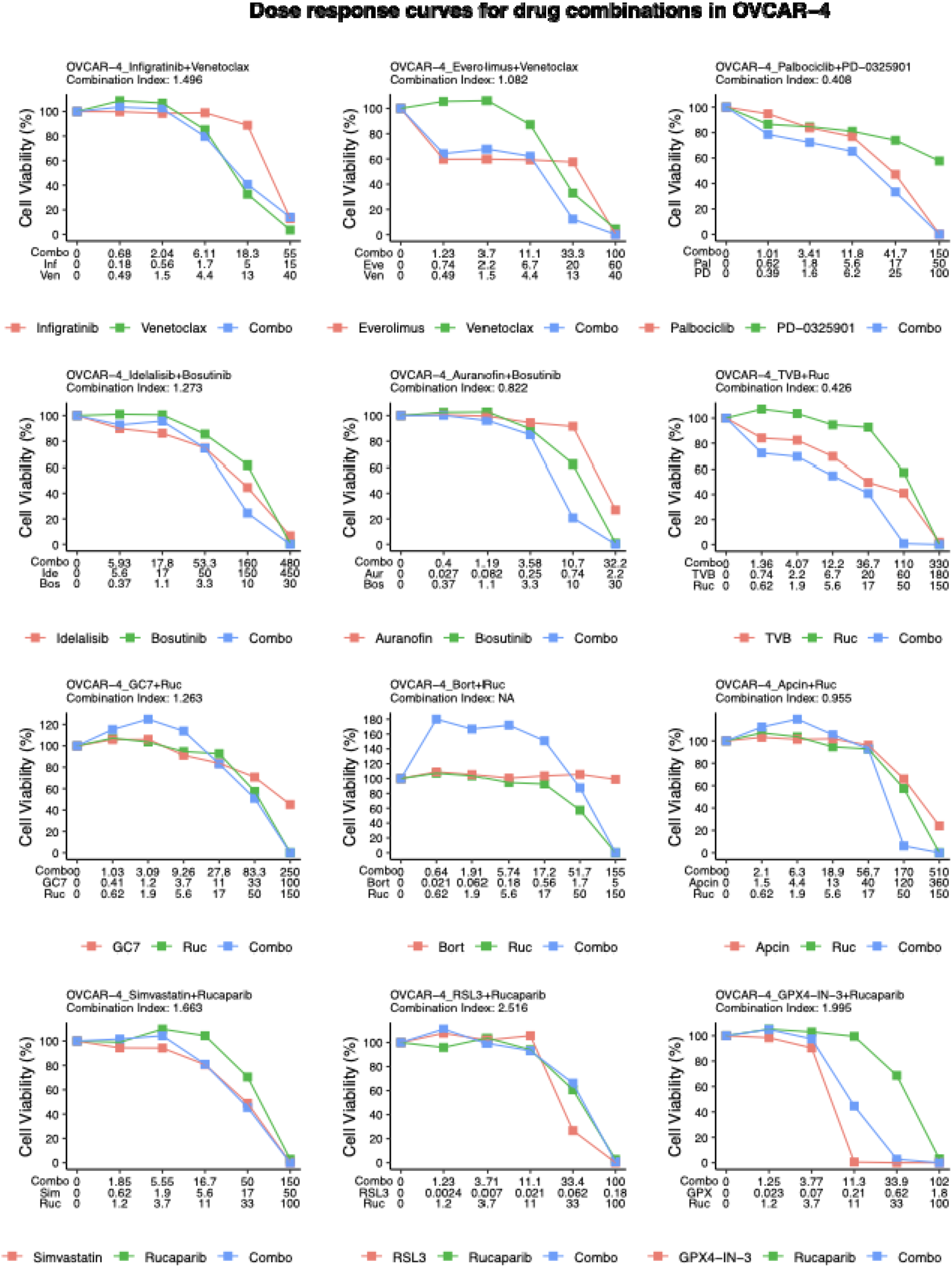
Inhibition of proliferation by various drug combinations in a representative ovarian cancer cell line (OVCAR-4). The combinations were chosen based on pathway analysis of protein response (Fig. 3). TVB-2640 (‘TVB’, FASNi), rucaparib (‘Ruc’, PARPi), bortezomib (‘Bort’, proteasome inhibitor), Apcin (APCi), GC7 (DNA synthase inhibitor), novobiocin (‘Novo’, DNA polymerase inhibitor), infigratinib (‘Inf’, FGFR3i), venetoclax (‘Ven’, BCL2i), everolimus (‘Eve’, MTORi), Palbociclib (‘Pal’, CDK4/6i), PD-0325901 (‘PD’, MEKi), idelalisib (‘Ide’, PI3Ki), bosutinib (‘Bos’, SRCi), auranofin (‘Aur’, PRDXi), simvastatin (‘Sim’, statin), GPX4-IN-3 (‘GPX’, GPX4i), RSL3 (GPX4i), DM4 (tubulin inhibitor). Each drug was applied to OVCAR-4 cells with a serial dilution factor of 3 or 4 to establish a dose-response curve for the single drug. Each drug combination was applied at the same concentrations as the single drugs. The combination index for each drug pair is calculated using the Chou-Talalay method.

#### Several combinations with MEKi and SRCi are anti-resistance candidates in ovarian cancer cells

The drug combination with synergy across most cell lines (based on CI) was the PD-0325901+Palbociclib drug combination targeting MEK and CDK4/6, respectively. This drug combination has been studied pre-clinically in colorectal cancer (C. L. Lee et al. 2023) and is the subject of an ongoing clinical trial for solid tumors (NCT02022982).

Several drug combinations, including those targeting NOTCH + AKT, MEK + EGFR, PKC + FASN, and those targeting IDH1 were only tested in the OVSAHO cell line due to resource limitations (STable 1). Drug combination with bosutinib (SRCi) had a synergistic effect in TOV-21G with idelalisib (PI3Ki), with a weak synergy effect in cell lines: IGROV-1 and OVCAR-4 (Table 2).

## Discussion

### Summary of the current study

There is great interest in rational approaches to the identification of drug combinations that can overcome initially present or acquired drug resistance in cancer. Adaptation to the stress of treatment can occur within a matter of days as cellular processes respond to support survival (Marine, Dawson, and Dawson 2020) (Bell et al. 2019). We conducted a comprehensive evaluation of protein level changes in high-grade serous ovarian cancer (HGSOC) using mass spectrometry (MS) to obtain insight into markers and pathways of resistance. In single-agent experiments using 7 drugs targeting key cancer processes (e.g., WNT, RAS/RAF/MEK, PI3K/AKT, apoptosis, etc.) in the OVSAHO cell line (chosen for its relative similarity to tumors), we identified responses in specific cellular processes using NetBox (see Methods) analysis that groups proteomic responses into functional modules. Suitable member proteins of a module were flagged as targets of a second agent in combination with the initial drug to prevent or overcome resistance. These hypotheses were followed up experimentally on a subset of several diverse cancer-derived cell lines that reasonably represent the diversity of HGSOC patient samples, as assessed by the similarity of genomic and expression profiles of the cell lines to those of HGSOC surgical samples (Table 3) (Domcke et al. 2013; Sinha et al. 2021).

Synergy is not an absolute requirement in the identification of useful drug combinations, nor a necessity for anti-resistance effect by the combinations. Additivity is sufficient as the second drug increases the anti-proliferative effect at full concentration. Synergy can be an advantage in terms of lower side effects by lowering the required concentration, but also a drawback as cancer cells adapt to perturbation: if sensitivity to one drug is lost, the synergistic effect is also lost (Palmer 2017). However, a number of the suggested and tested drug combinations are not suitable for further investigation, as their anti-proliferative effect is substantially antagonistic (CI > 2.0, Table 2).

### Response to PARPi combinations in genetically diverse cell lines

This workflow revealed several effective drug combinations. Building on our previous work on PARP inhibitors, we see that PARPi+FASNi has a strong anti-proliferative effect versus single-agent PARPi in OVCAR8 cells, which aligns with a growing interest in the importance of lipid metabolism in ovarian cancer and possible therapeutic implications (Yoon and Lee 2022; Chaudhry, Thomas, and Simmons 2022; Ji et al. 2020). Related to lipid requirements of cell proliferation, we did explore the combination of PARPi (rucaparib) with statins (i.e., simvastatin), which target HMG-CoA reductase, a key enzyme in the mevalonate pathway. With this combination, we also see a synergistic effect in the OVCAR8 cell line. The unique genomic profile of the OVCAR8 may contribute to these observations. OVCAR8 has an AA-changing mutation (V613L; variant of unknown significance, VUS) in the ATM gene that is related to homologous recombination (HR) and has no other HR-related mutations (Table 3). Unique to OVCAR8 is a KRAS P121H in a loop near the G base of GTP, which may affect GDP/GTP exchange. We do see some overlap in drug-drug synergy patterns between the OVCAR8 and the OVCAR4 cell lines; we see synergy involving PARPi+APCi in both OVCAR8 and OVCAR4. Both cell lines have been reported by several groups as HR proficient, and both have an ATM mutation (N230T in OVCAR4, a VUS) (Wilson et al. 2018; Kondrashova et al. 2018; Siddiqui et al. 2021; Lu et al. 2022). The results using combinations with PARPi build on our previous discussion on the therapeutic benefit of PARP inhibition regardless of HR status (Franz et al. 2021; González-Martín et al. 2019).

### Ongoing interest in the inhibition of GPX4

Regarding HR deficiency, recent work has shown that BRCA1-deficiency may be important to the response of cancer cells co-treated with a GPX4 and PARP combination, where GPX4 inhibition can induce ferroptosis (Lei et al. 2024). Of the cell lines we selected, only IGROV-1 is BRCA1-deficient; therefore, it is difficult to draw conclusive direct comparisons. Of the ovarian cell lines tested in this study, IGROV1, OVCAR4, and OVSAHO are BRCA2-deficient that could affect response. In related work, in HT1080 fibrosarcoma BRCA2-knockout cells, Lei et al. do not observe a difference in RSL3 response (Lei et al. 2024). Across the cell lines tested, we see no synergy as measured by the combination index with a GPX4i and PARPi combination (Table 2). That said, our treated cell lines were highly responsive to both of the tested GPX4 inhibitors as single agents (RSL3 and GPX4-IN-3; IC50 0.5 uM and 0.11 uM), suggesting further preclinical work. One issue hampering the further development of RSL3 for clinical application is its reported low aqueous solubility (Gaschler et al. 2018; L. Wang, Chen, and Yan 2022). There is continuing research to develop novel GPX4 inhibitor chemistry, as well as means of targeted delivery to overcome existing limitations (Eaton et al. 2020; W. Li et al. 2022; Gao et al. 2019). Separately, other groups are exploring other novel combinations utilizing GPX4 inhibition. NRF2/GPX4 (RSL3) combinations in ovarian cancer and head and neck cancer (N. Li et al. 2024; Shin et al. 2018) and Taxol in GPX4 inhibit the proliferation of cell lines (Feng et al. 2023). The rationale for the NRF2/GPX4 combination is that NRF2 is a regulator of anti-ferroptic genes, including genes that prevent the accumulation of free iron (Dodson, Castro-Portuguez, and Zhang 2019). This work with GPX4 is of particular interest, as there have been observations of resistance to platinum-based treatments and GPX4 levels in ovarian patients (X. Wu et al. 2022). So both single-agent inhibition of GPX4 and combinations may be particularly promising for pre-clinical studies in ovarian cancer.

### Further proposed pre-clinical work

For the successful combinations and single agent result we describe here, further study in more advanced preclinical models (e.g., organoids, patient-derived xenografts (PDXs), immune-competent mouse models), rather than in 2D cell lines as done here, is warranted, given potential insights that could be obtained, but outside of the scope of the current work (Dorrigiv et al. 2021; Kapałczyńska et al. 2018; Jensen and Teng 2020; Imamura et al. 2015). If conducted, such studies could provide an initial assessment of toxicity for the proposed drug combinations, as well as some understanding of the effectiveness of the treatment in the presence of immune cells. To our knowledge, none of the proposed drug combinations have been subject to clinical trials specific to ovarian cancer. In future work, we and others are encouraged to test the proposed combinations in organoids, PDXs, and animal models. Another proposal for future study is to explore staggered rather than simultaneous exposure to two drugs. In the experiments reported here, cells were simultaneously treated with a given drug combination. Staggered (or sequential) treatment in preclinical studies is less common. Several preclinical studies demonstrate the efficacy of a staggered approach and propose mechanistic rationales (Settleman, Neto, and Bernards 2021; M. J. Lee et al. 2012). Specific to PARPi treatments, there is some evidence for the utility of PARPi in combination with other treatments (BETi and WEEi) under a given sequential regimen (Fang et al. 2019; Peng et al. 2024). There is a need for computationally reliable predictive techniques that optimize sequential multi-drug treatments based on time-dependent (dynamic) models.

### Possibilities for future developments at larger scales

In previous work, we developed a perturbation biology framework (CellBox) (Yuan et al. 2021) for the derivation of computational models predictive of the cellular response to perturbations of any one of the many proteins observed in perturbation-response experiments, or even of combinations of proteins. Such models would be potentially more powerful and replace the module-plus-target-identification analysis used here, but their parameterization requires a much larger number of systematic perturbation experiments: probably hundreds or thousands of targeted perturbations, many more than the seven used here. This scale-up remains an open challenge and an opportunity, provided by the scale of thousands of protein levels quantitatively measurable by mass spectrometry. The work presented here provides guidance to the design of ongoing efforts that make use of comprehensive protein and phosphoprotein changes resulting from the perturbation of key cancer processes (Kupcik et al. 2019)(J. Li et al. 2022; Franciosa et al. 2023)(J. Li et al. 2022).

## Materials and Methods

### Drug perturbation experiments for proteomic profiling

OVSAHO cells were obtained from JCRB (Japanese Collection of Research Bioresources Cell Bank) cell bank (NIBIOHN, Cell No. JCRB 1046) and grown in MCDB105/199 medium supplemented with 10% FBS (Fisher, #10438026) and 1% Penicillin-Streptomycin. All cells were free of Mycoplasma, and their identity was verified by whole-exome sequencing at the sequencing platform of the Broad Institute of MIT and Harvard. The following drugs used in the experiments were all commercially available: AKT inhibitor MK-2206 2HCl (Selleckchem, #S1078), BCL2 inhibitor Venetoclax (ABT-199, Selleckchem, #S8048), GSK3β inhibitor CHIR-99021 (Selleckchem, #S1263), MEK inhibitor PD-0325901 (Selleckchem, #S1036), PKC inhibitor Bisindolylmaleimide VIII (Caymanchem, #13333), and SRC inhibitor Bosutinib (SKI-606, Selleckchem, #S1014). To determine the IC50 for each of the drugs, cells were treated with DMSO control or varying concentrations (from 0.01 μM to 100 μM) of each drug 24 hours after seeding. The Incucyte NucLight Rapid Red (NRR) Reagent (Essen Bioscience, #4717, 1:4000) was added to the medium to label the nuclei of live cells. Cell viability was determined as the number of live cells 72 hr after drug treatment, counted using the live imaging of Incucyte, normalized to those of the DMSO control. Cell viability data from three biological replicates, each consisting of three technical replicates, were merged and fitted using a non-linear log(inhibitor) versus normalized response with a variable slope for dose-response curves using GraphPad Prism. IC50 was determined as the drug concentration that gives 50% of the maximal inhibitory response on cell viability for each drug. To harvest cells for mass spectrometry proteomic profiling, cells were treated with DMSO control or the IC50 concentration 24 hr after seeding in 10 cm dishes for each drug treatment. Cells were harvested 72 hours after drug treatment by gently scraping off from the plate into PBS and transferred into deep 2 mL 96-well plates. Cells were stored at -80°C and sent for mass spectrometry measurements.

### Liquid chromatography-mass spectrometry (LC-MS) measurements and MS data analysis

Harvested cells were digested, and the proteomic measurements were taken as previously described (Franz et al. 2021), section ‘Liquid chromatograph-mass spectrometry (LC-MS) measurements’ of Methods. The raw data obtained from mass spectrometry measurements were processed as previously described (Franz et al. 2021), section ‘MS data analysis’ of Methods. The protein expression matrix with rows as identified proteins and columns as samples was obtained after data processing.

### Proteomics data analysis

The protein expression matrix contained protein measurements from three biological replicates of each drug treatment and two sets of three biological replicates of DMSO control (due to two batches of samples). Data analysis was performed using customized scripts in R (Version 3.6). Counting the number of measured proteins, multidimensional scaling (MDS) on Euclidean distance of protein measurements, and pairwise Pearson correlation of individual samples were performed on the raw protein expression matrix. To identify differentially expressed proteins of each perturbation condition from negative controls, an unpaired t-test was used to compare protein expressions from the drug-treated samples with samples treated with DMSO. log2(expression ratio) (‘log2ratio’) and corrected p-values for an FDR < 0.2 by the Benjamini-Hochberg (BH) method were obtained for each drug perturbation. Strongly responsive proteins were defined as those whose log2ratio > 0.5 or < -0.5 with corrected p-value < 0.05. The protein measurements from the biological replicates were then pooled to obtain average protein expression values for each perturbation condition. The averaged protein expression across replicates was then used for unsupervised hierarchical clustering, pairwise correlation of perturbation conditions, and all the following analyses. Visualization using the upset plot was performed with UpSetR (Conway, Lex, and Gehlenborg 2017).

### Protein module detection and enrichment analysis

NetBox was used to perform the detection of responsive protein modules (Cerami et al. 2010; Liu et al. 2020). For each of the drug-treatment conditions, the input gene list contains strongly responsive proteins upon the drug treatment (absolute value of log2ratio > 0.5) whose p-value of the t-test of differential expression is smaller than 0.01. To ensure the most compact and confident module findings, the adjusted p-value of the connected linker node was set to 0.005, with all the other parameters of NetBox kept at default. Two protein-protein interaction networks were used independently as the background network of NetBox analysis: 1) Reactome Functional Interaction (FI) Network (version 2020 from Reactome.org) (Gillespie et al. 2022) with predicted interactions filtered out, 2) Interactions collected by Integrated Network and Dynamical Reasoning Assembler (INDRA) with at least a belief score of 0.95 (Gyori et al. 2017), which typically indicate the interaction is supported by multiple independent reading systems or pathway databases. Enrichment analysis was performed on the identified modules using the clusterProfiler Bioconductor package (Yu et al. 2012) using gene annotations from the Gene Ontology with default parameters. Gene Set Enrichment Analysis (GSEA) (Subramanian et al. 2005) was performed on the responsive proteins using clusterProfiler with default parameters. The input gene lists for each drug treatment are consistent with those for NetBox analysis, and the ranking of the genes is based on the log2ratio of the corresponding proteins in decreasing order.

### Determination of functional score

The list of cancer census genes was obtained from the Catalogue of Somatic Mutations In Cancer (COSMIC, cancer.sanger.ac.uk) (Tate et al. 2019). The functional score of a gene is defined as positive (1, pro-proliferative) if the role of the gene in cancer is an oncogene, and negative (-1, anti-proliferative) if the role of the gene is a tumor suppressor gene (TSG). If a gene has both roles of oncogene and TSG, the functional score is not defined.

### Combination perturbation experiments for candidate validation

Experimental validation of the proposed combination candidates was performed by Charles River Laboratories. Six cell lines were used in the experiments, including IGROV-1 (NCI #0507369, RRID: CVCL_1304), OVCAR-4 (NCI #0502527, RRID:CVCL_1627), OVCAR-5 (NCI #0507336, RRID:CVCL_1628), OVCAR-8 (NCI #0507407, RRID:CVCL_1629), OVSAHO (RRID:CVCL_3114), and TOV-21G (ATCC CRL-11730 Lot#58690706, RRID:CVCL_3613). OVSAHO cells were grown in MCDB105/199 medium supplemented with 10% FBS and 1% Penicillin-Streptomycin. TOV-21G cells were grown in RPMI medium supplemented with 15% FBS and 1% Penicillin-Streptomycin. All other cell lines were grown in RPMI-1640 medium supplemented with 10% FBS and 1% Penicillin-Streptomycin. All the small-molecule drugs are commercially available: Rucaparib (Selleckchem, #S1098), TVB-2640 (Selleckchem, #S9714), Bortezomib (Selleckchem, #S1013), Apcin (Selleckchem, #S9605), GC7 (Sigma-Aldrich, #259545), AL101/BMS-906024 (MedChemExpress, #HY-15670), Infigratinib (Selleckchem, #S2183), Everolimus (Selleckchem, #S1120), Ivosidenib (Selleckchem, #S8206), Erlotinib (Selleckchem, #S7786), Palbociclib (Selleckchem, #S1116), Bosutinib (Selleckchem, #S1014), Idelalisib (Selleckchem, #S2226), and Auranofin (Selleckchem, #S4307).

Cells were seeded into 96-well plates for checkerboard assays, in which a 3-fold serial dilution of one compound in the combination candidate was performed vertically down the plate for 6 dilution points, and a 3-fold serial dilution of the other compound was performed horizontally across the plate for 9 dilution points. One column of DMSO controls was included for each plate. A 3-fold serial dilution of 6 dilution points was also performed for each compound in single-drug treatment experiments. Different concentration ranges were tested for different compounds for the best outcome: Rucaparib (6 3-fold dilution from 150 μM to 0.62 μM), TVB-2640 (9 3-fold dilution from 180 μM to 0.027 μM), Bortezomib (9 3-fold dilution from 210 nM to 0.032 nM), Apcin and GC7 (9 3-fold dilution from 360 μM to 0.055 μM). Cell viability was measured using the CellTiter-Glo assay 72 hr after drug treatment. Synergy scores were calculated using the SynergyFinder R package (Zheng et al. 2022).

## Author Contribution

AF, CiSh, FC, MM, CS were involved in the conceptualization of the study; AF, CiSh, FC, MM worked on collecting the data; AF, CiSh, KM, LC worked to perform investigative experiments; AF, CiSh, AL wrote and ran analysis scripts; AL, CS, MM carried out various supervisory functions; AF, CiSh, AL, AK, CS analyzed the data. AF, CiSh, AL, CS wrote the original draft along with the original figures; all authors reviewed and edited the final manuscript.

## Availability of data and materials

Processed data is provided within the manuscript or supplementary information files. The mass spectrometry proteomics data have been deposited to the ProteomeXchange Consortium via PRIDE with accession number PXD066316.

## Declarations

### Ethics approval and consent to participate

Not applicable.

### Consent for publication

Not applicable.

## Supporting information

SFigure 3

STable 2

STable 1

STable 3

