## Supplementary material for "Design of combination therapeutics from protein response to drugs in ovarian cancer cells": SFigure 3

### Dose response curves for drug combinations in IGROV−1

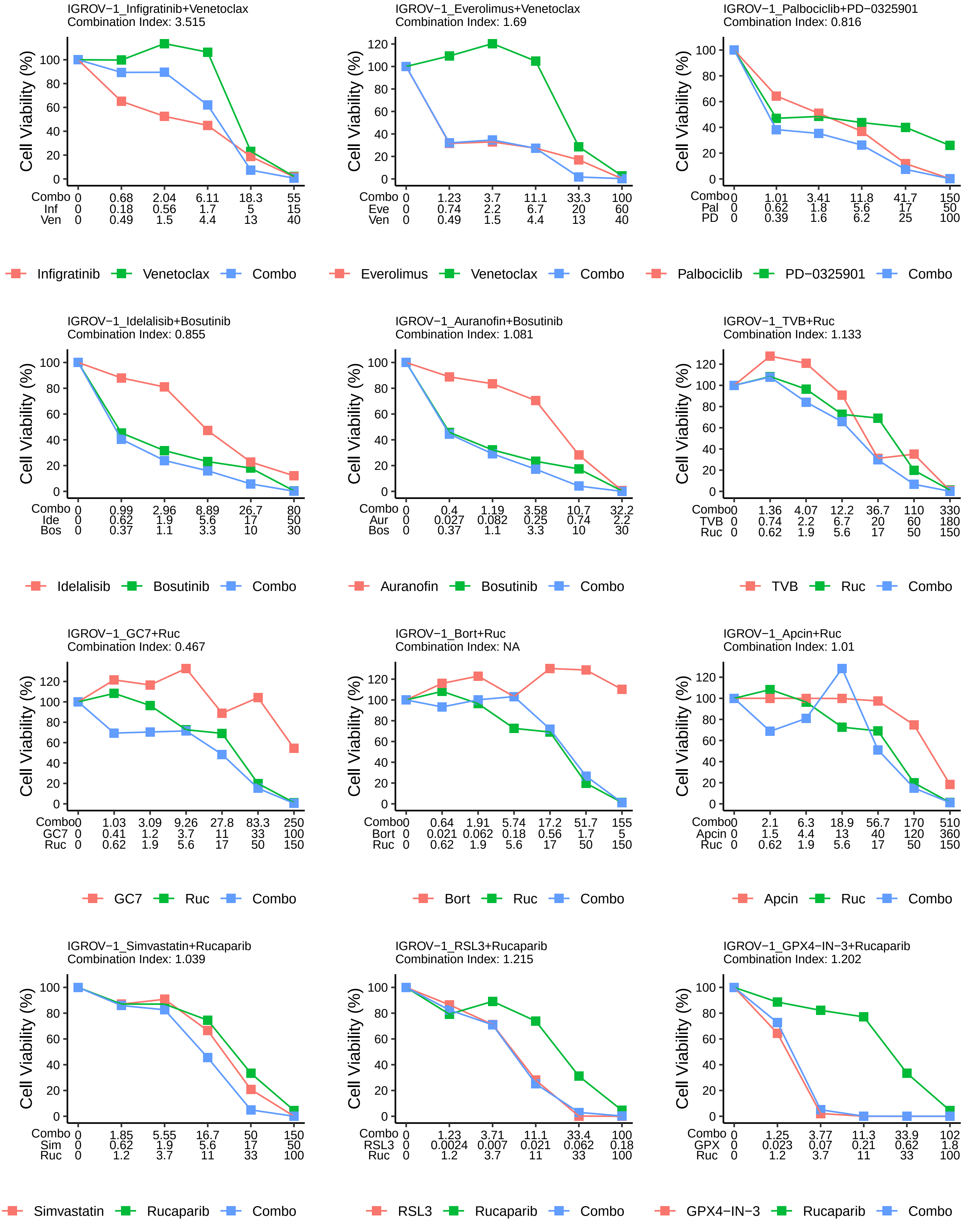

### Dose response curves for drug combinations in OVCAR−4

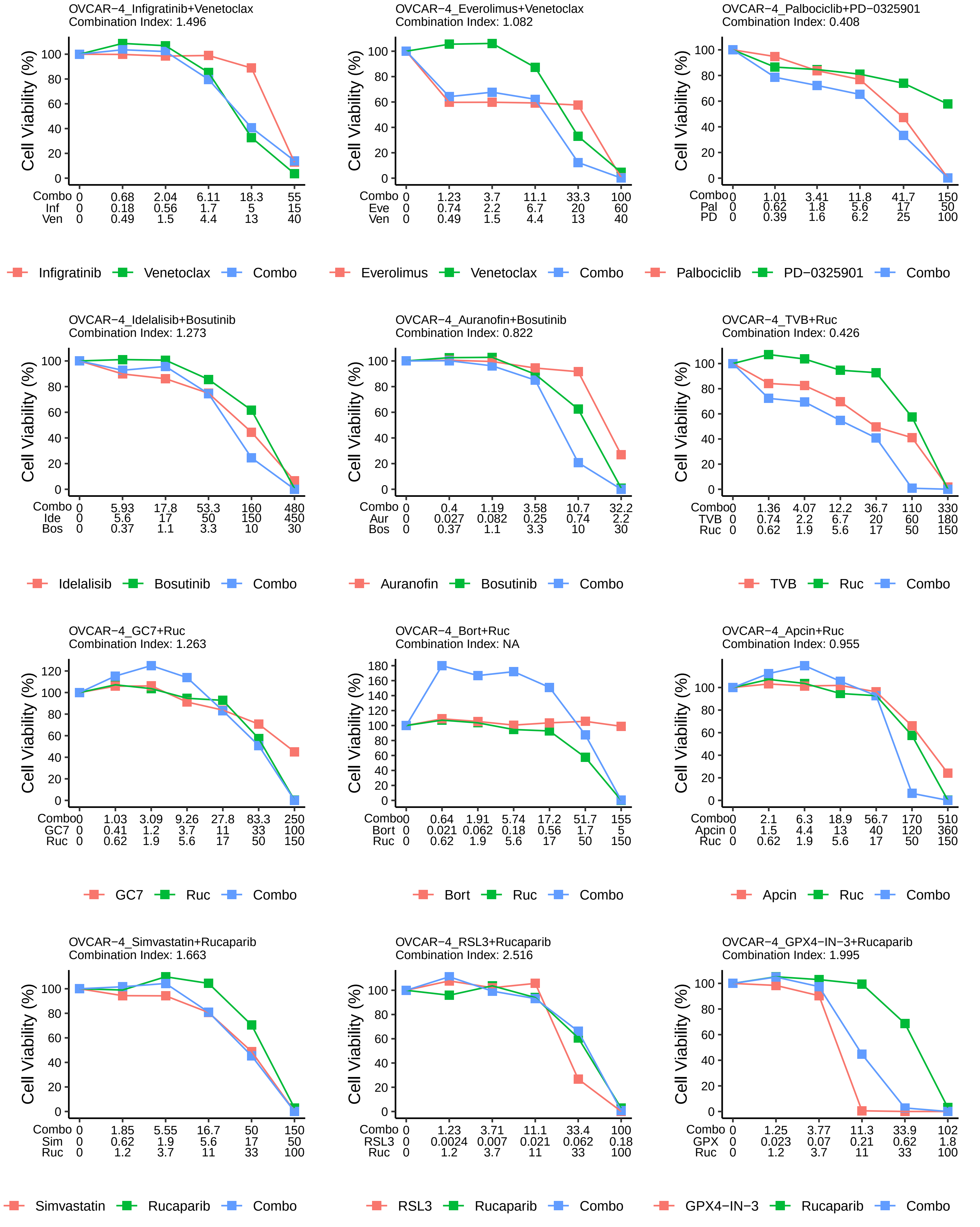

### Dose response curves for drug combinations in OVCAR−5

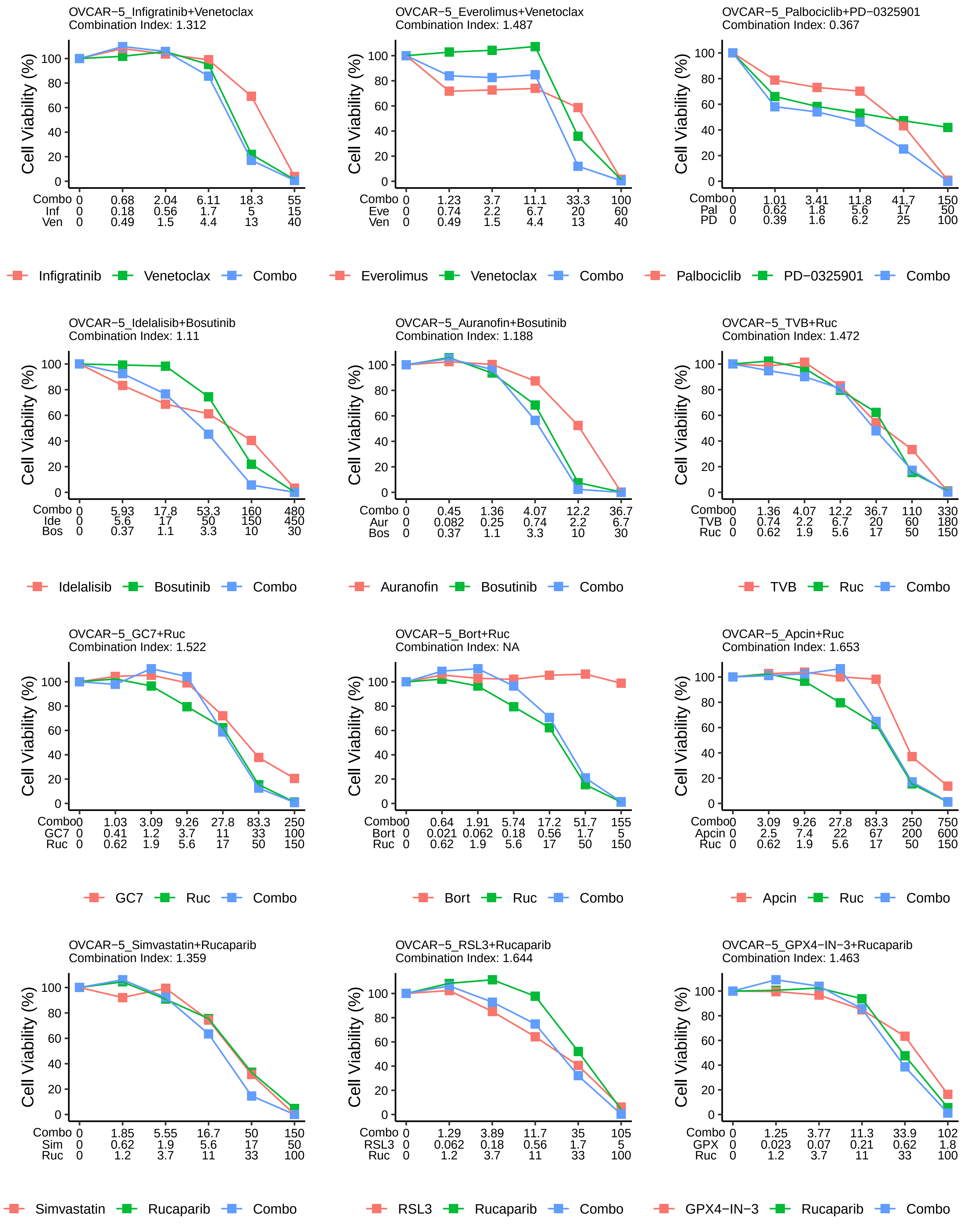

### Dose response curves for drug combinations in OVCAR−8

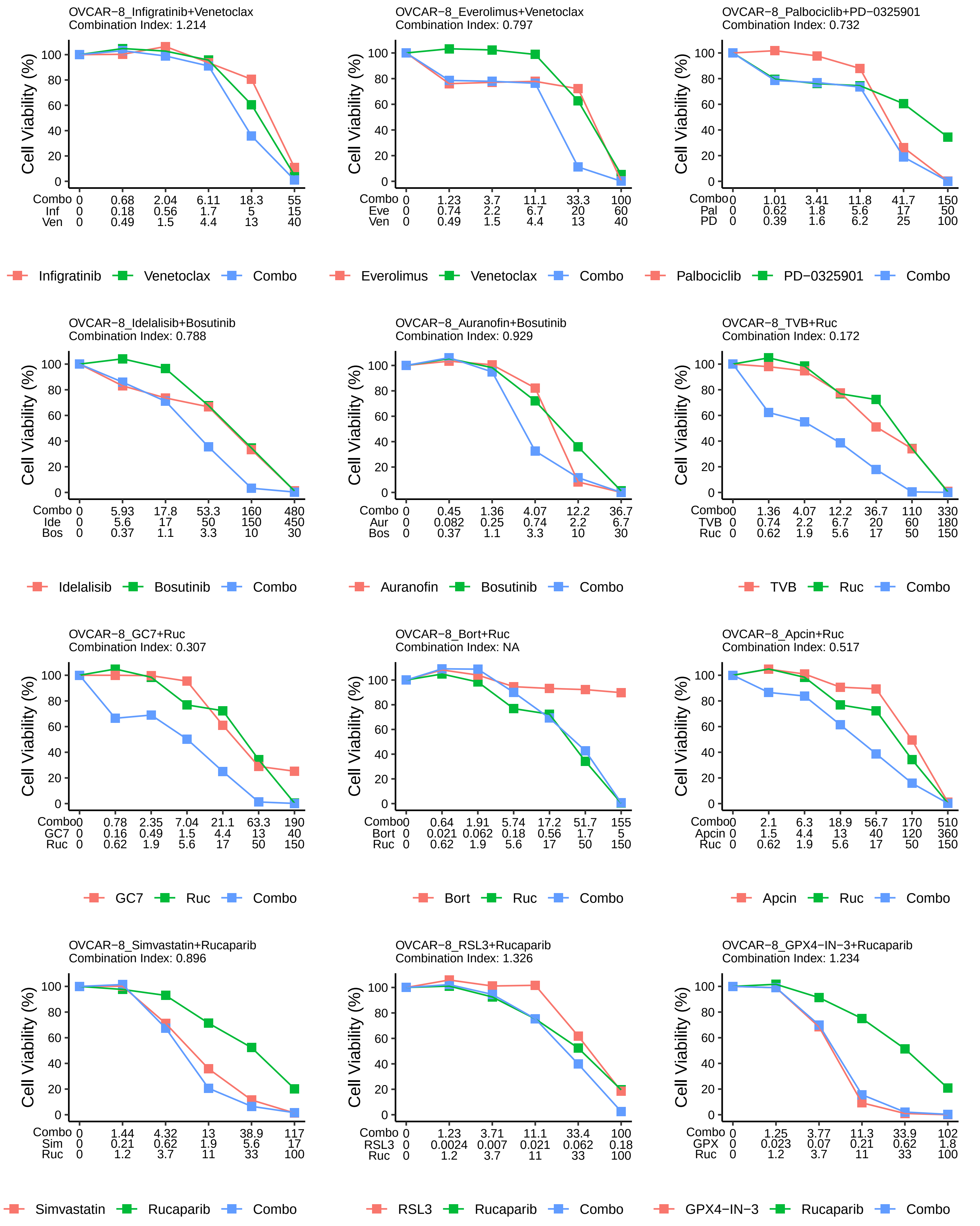

### Dose response curves for drug combinations in OVSAHO

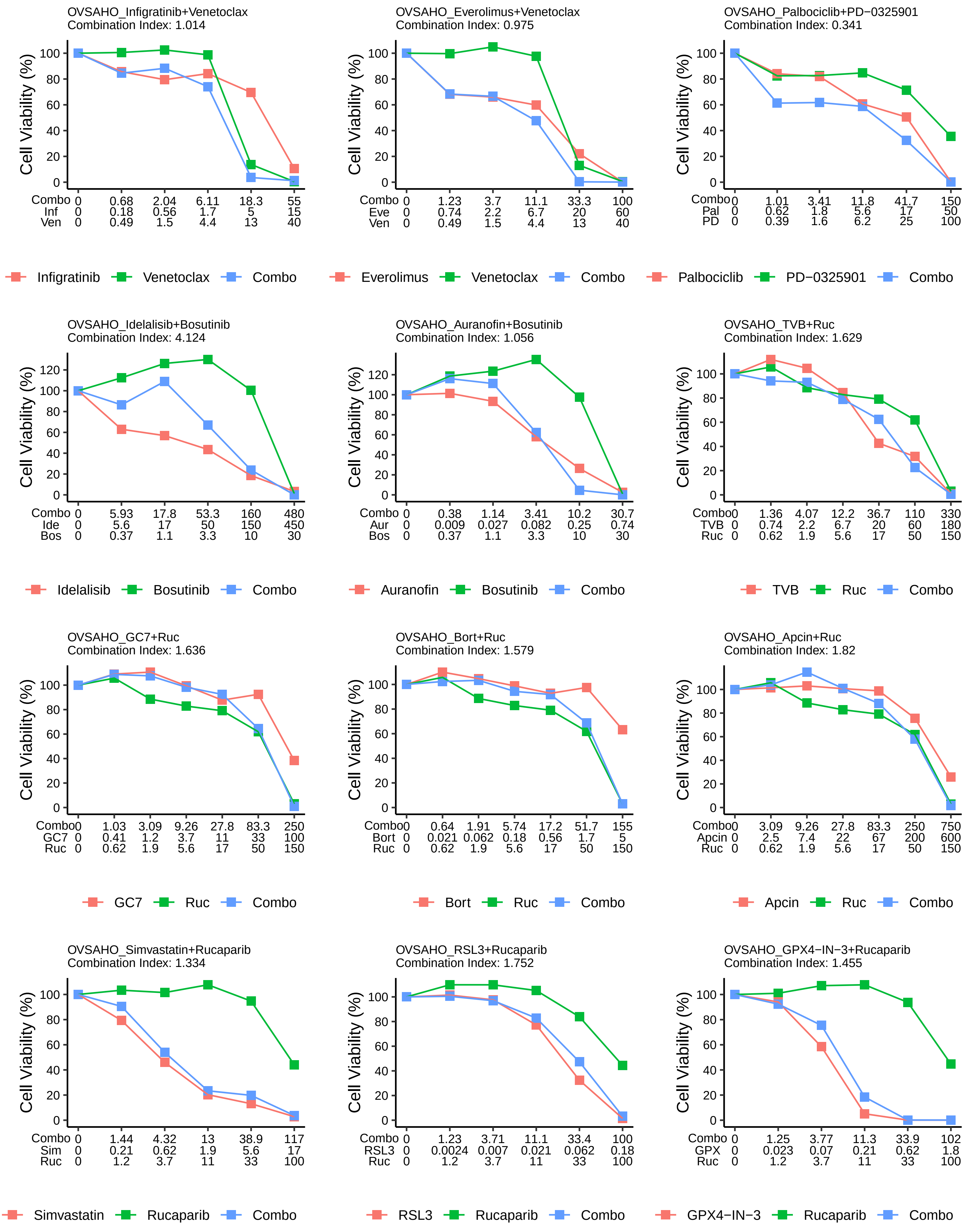

### Dose response curves for drug combinations in TOV−21G

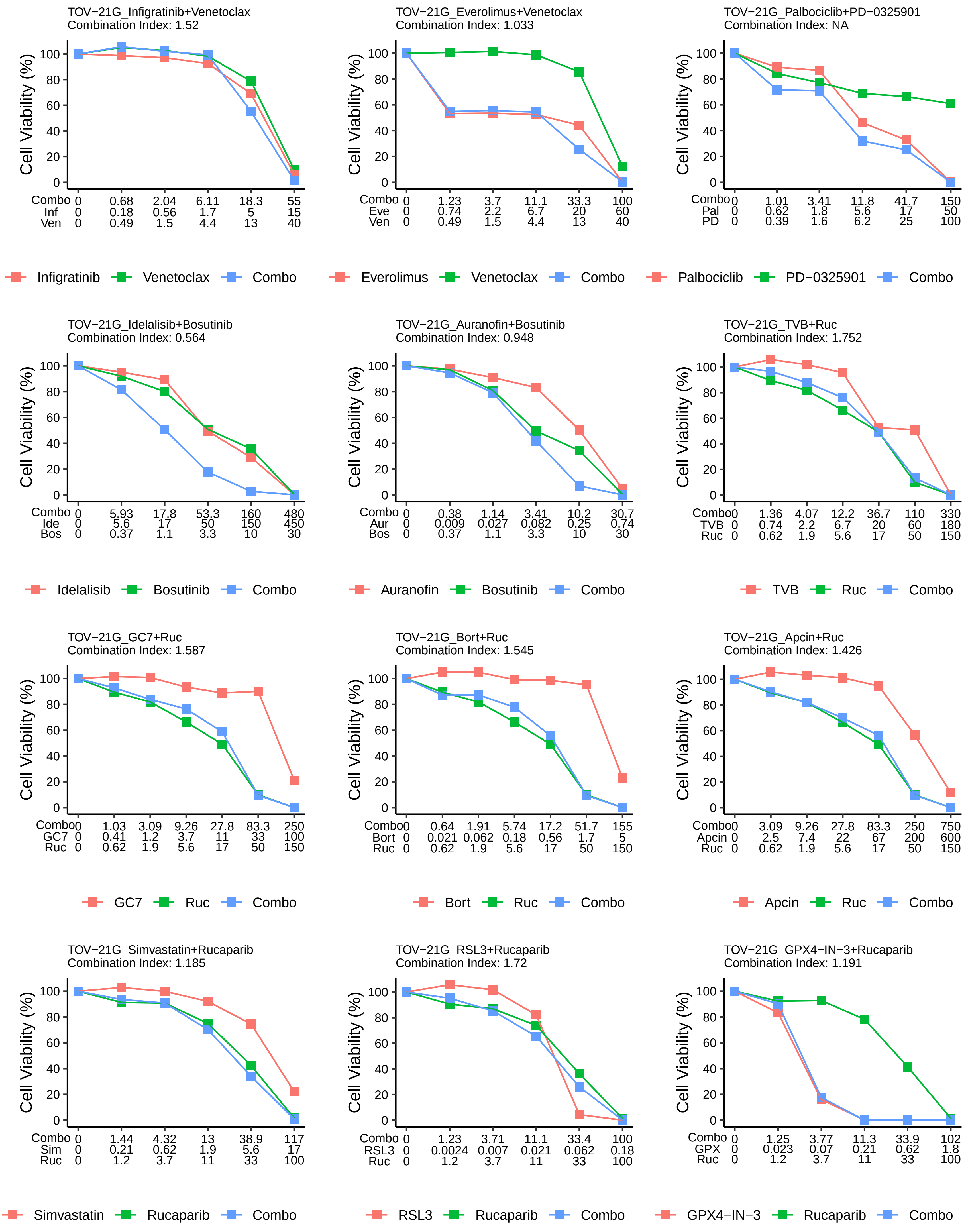
