## Supplementary notes for "Design of combination therapeutics from protein response to drugs in ovarian cancer cells"

### SUPPLEMENTARY MATERIALS

#### Supplementary notes

For each of the small-molecule drugs, the protein modules with decreased activities identified using Netbox analysis, indicated by the decrease in protein expressions, generally agree with the known functions of the drug. The full list of identified modules using both the FI and INDRA networks is shown in supplementary table STable 3.

AKTi In agreement with AKT's central function in controlling key kinase signaling pathways, cell proliferation, and glucose metabolism (Manning and Toker 2017; Nitulescu et al. 2018), we found that the responsive protein network modules upon AKT inhibition are enriched for several biological pathways, including transmembrane receptor protein serine/threonine kinase signaling (GO:0007178, adj. p-value = 0.0024), muscle contraction (GO:0006936, adj. p-value = 2.10E-05), nuclear chromosome segregation (GO:0098813, adj. p-value = 0), regulation of fibroblast proliferation (GO:0048145, adj. p-value = 0.044). The decrease of activities in these biological processes, indicated by the decrease in protein expression, likely led to the decrease in cell viability.

BCLi By analyzing the protein network modules derived upon BCL2 inhibition, we found that BCL2i negatively affected proteins involved in mitochondrial translation (GO:0032543, adj. p-value = 0), and protein import into the mitochondrial matrix (GO:0030150, adj. p-value = 0). This is in agreement with the role of BCL2 in mitochondrial apoptosis and metabolism (Giménez-Cassina and Danial 2015).

GSK3βi Inhibition of GSK3β led to the disrupted activities of several biological pathways, including Golgi vesicle transport (GO:0048193, adj. p-value = 0), and regulation of canonical Wnt signaling pathway (GO:0060828, adj. p-value = 0.0060), which are consistent with the known functions of GSK3β to regulate Golgi membrane trafficking (Adachi et al. 2010) and affect Wnt/β-catenin signaling (Huang et al. 2017).

MEKi We found that the responsive protein modules upon MEK inhibition are enriched for biological processes involved in cell cycle and proliferation, including mitotic sister chromatid segregation (GO:0000070, adj. p-value = 0), mitotic spindle assembly (GO:0090307, adj. p-value = 2.17E-05), and ribosome biogenesis (GO:0006364, adj. p-value = 0). These observations are consistent with the major function of the MAPK/ERK signaling pathway to regulate cell cycle entry, transcription, and translation.

PKCi Enriched biological processes upon PKC inhibition include extracellular matrix organization (GO:0030198, adj. p-value = 0), establishment of vesicle localization (GO:0051650, adj. p-value = 3.17E-05), regulation of small GTPase mediated signal transduction (GO:0051056, adj. p-value = 1.54E-06), etc. Altered activities in these processes agree with known regulatory functions of PKC, including but not limited to membrane structure, vesicle permeability, and phosphorylation of downstream proteins.

SRCi Protein modules identified upon SRC inhibition are enriched mainly for biological processes that are essential for cell survival and proliferation, including covalent chromatin modification (GO:0016569, adj. p-value = 0), transcription elongation from RNA polymerase II promoter (GO:0006368, adj. p-value = 0), etc.

**SFigure 1**

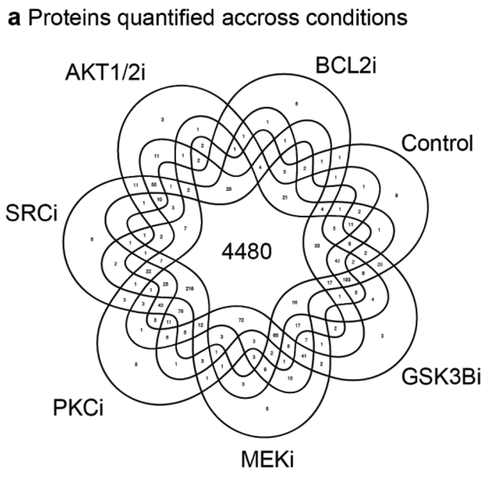


(a) Venn diagram illustrating the overlap of proteins captured and quantified across treatment conditions with a total of 5912 proteins identified, and 4480 proteins found in all perturbation conditions.

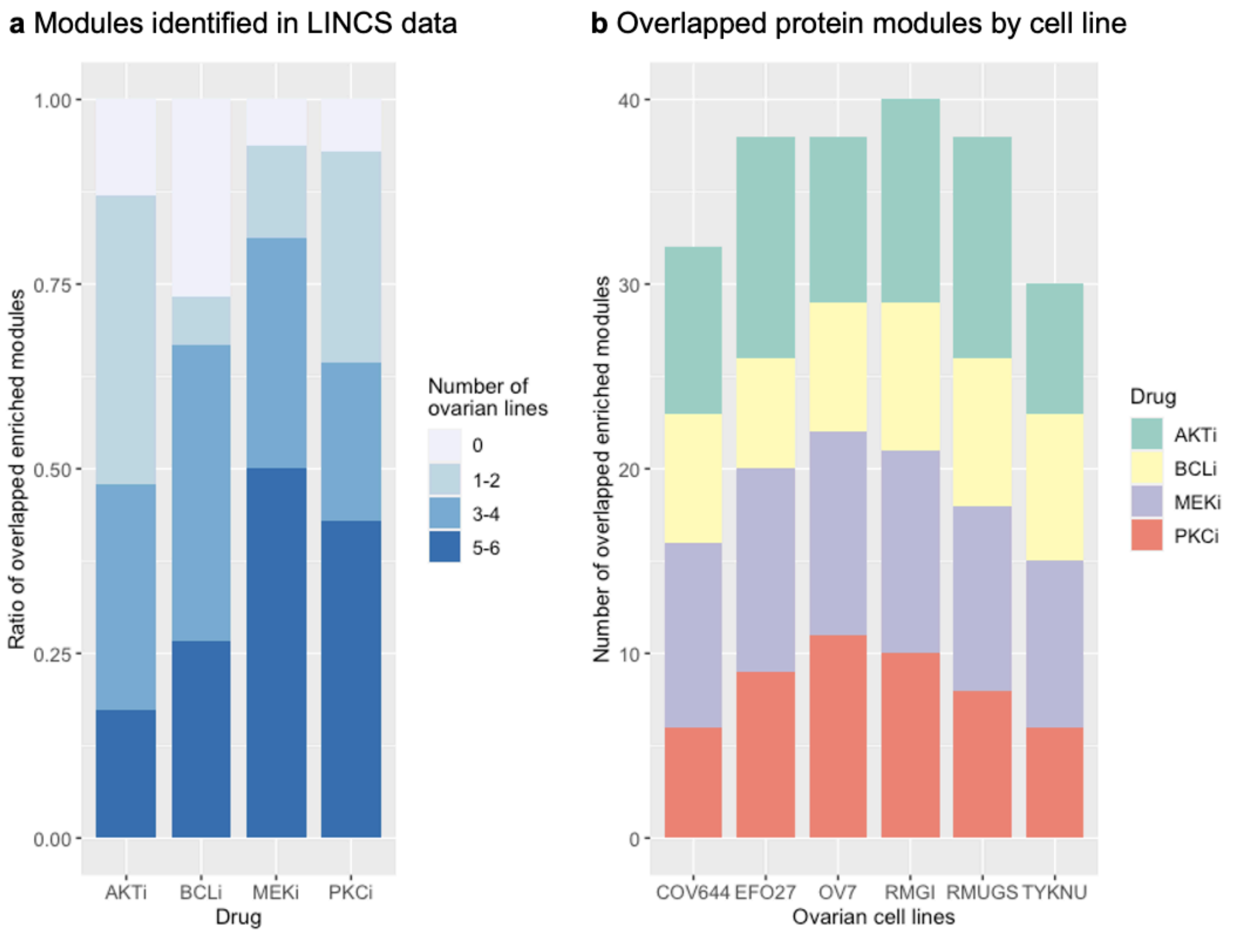


**SFigure 2**

**Identified protein modules validated in an independent dataset.** (a) Protein module identification by NetBoxR and enrichment analysis have been performed on the Library of Integrated Network-Based Cellular Signatures (LINCS) perturbation-response dataset. Most of the identified modules (>70%) in mass-spec data can be identified in at least one of the six ovarian cell lines in LINCS data. Many modules (>50%) can be identified in more than two cell lines. (b) Each of the ovarian cell lines has a significant number of identified protein modules overlapped with those identified in OVSAHO, while RMGI, which is the most similar one to OVSAHO in terms of HGSOC properties, has the highest number of overlapped modules.

**SFigure 3**

**dose_response_curves_combined.pdf:** Dose-response curves for single drug and combination pairs tested in 6 ovarian cell lines

**STable 1**

**ic50_values_used_for_proteomic_profiling.csv:** IC50 of individual drugs on OVSAHO cell line derived from dose-response curves. These drug concentrations were used for the proteomic profiling experiments.

**STable 2**

**drugs_dataset_median_corrected_NA_ratio.csv:** Protein expression levels across different perturbations. The data table includes raw protein expression (column labeled as perturbation drug name_replicate), averaged (mean_), standard deviation (SD_) and variation (CV_) of protein expression across replicates, ratio of perturbed over control (DMSO) (ratio_) and p-value from t-test (p.value_). The data table was used for all analyses in this study.

**STable 3**

**single_drug_ic50_summary.csv:** Single drug IC50 values tested across 6 ovarian cell lines. IC50 of all the drugs involved in the combination candidates tested in this study. These IC50 values were used to compute combination indices CI for the drug combinations.
